# Sex differences in astrocytic reactivity and its association with cognitive decline in cognitively unimpaired older adults

**DOI:** 10.64898/2026.01.30.702902

**Authors:** Chelsea Reichert Plaska, Tovia Jacobs, Davide Bruno, Sang Han Lee, Bruno P. Imbimbo, Ricardo S. Osorio, Andréa L. Benedet, Nicholas J. Ashton, Henrik Zetterberg, Nunzio Pomara, the Alzheimer’s Disease Neuroimaging Initiative

**Author notes:** Corresponding Author: Nunzio Pomara, MD Nathan S. Kline Institute for Psychiatric Research 140 Old Orangeburg Road, Bldg. 35 Orangeburg NY 10962. Data used in preparation of this article were obtained from the Alzheimer’s Disease Neuroimaging Initiative (ADNI) database (adni.loni.usc.edu). As such, the investigators within the ADNI contributed to the design and implementation of ADNI and/or provided data but did not participate in analysis or writing of this report. A complete listing of ADNI investigators can be found at: http://adni.loni.usc.edu/wp-content/uploads/how_to_apply/ADNI_Acknowledgement_List.

## Abstract

**INTRODUCTION:** Astrocyte reactivity may contribute to increased Alzheimer’s disease (AD) risk in women. Although plasma glial fibrillary acidic protein (GFAP) levels have been reported to be higher in women than in men, few studies have examined sex-specific associations with other plasma AD biomarkers and cognition.

**METHODS:** Participants were enrolled in the Memory Education and Research Initiative, a longitudinal community-based cohort of individuals referred for memory evaluation. The Alzheimer’s Disease Neuroimaging Initiative (ADNI), a research-recruited cohort, was used for replication.

**RESULTS:** Females had higher plasma GFAP levels than males in MERI (N=584), a finding that was replicated in ADNI. In MERI, higher baseline GFAP was associated with lower Aβ42/40 and higher p-Tau231 and NfL levels. Higher baseline GFAP was also associated with steeper longitudinal decline in global cognition and episodic memory; however, GFAP × sex × time interactions were not significant. In ADNI, higher baseline GFAP was associated with higher p-Tau217 and NfL levels and with steeper longitudinal decline in global cognition and selected executive and semantic measures, without consistent evidence of sex-specific longitudinal effects. DISCUSSION: Plasma GFAP levels were consistently higher in females across two independent cohorts. GFAP was associated with AD-related blood biomarkers and longitudinal cognitive decline, but sex did not consistently modify these associations. These findings support a role for astrocytic reactivity in early AD-related processes while underscoring the need for further replication of sex-specific effects.

## 1 INTRODUCTION

Women account for two-thirds of all Alzheimer’s disease (AD) diagnoses^1^. Longer life expectancy contributes to AD prevalence^2,3^, but does not fully explain it, pointing to additional biological vulnerabilities. Additional risk factors include menopausal estrogen decline^4^, higher prevalence of affective disorders^5^, and a greater burden of inflammatory conditions^6^. Yet, despite increasing attention to sex differences in AD, the impact of sex on neuroinflammatory processes and their relationship to AD pathology remains poorly understood^7,8^.

Astrocytes are one of the key mediators of neuroinflammation^9–17^. In response to AD pathology, astrocytes become reactive, with increased expression of cytoskeletal glial fibrillary acidic protein (GFAP)^18,19^. Several lines of evidence, including post-mortem studies, have established elevated GFAP level in mild cognitive impairment (MCI) and AD, and its associations with greater AD pathology and cognitive decline^20–22^. Elevated GFAP levels have also been associated with increased future AD risk^23^ and predict progression to AD up to 15 years before clinical diagnosis^24^. While the aforementioned studies included sex as a covariate, they did not test whether sex modifies the association of GFAP with AD biomarkers – this approach minimizes or removes sex-related effects, rather than examining them as a meaningful biological variable^3,25^

Astrocytes are essential for supporting cognitive processes^26,27^. In cognitively healthy older adults, elevated GFAP levels are associated with poorer cognitive performance, particularly lower episodic memory scores^26,28^. Higher GFAP levels are also linked to worse performance on executive function tasks^29,30^ and reduced language abilities^28,30^. However, findings are not consistent; in at least one study comparing AD, other dementias and cognitively unimpaired (CU), GFAP showed no relationship with cognition^31^. Collectively, these findings highlight astrocytes as key contributors to cognitive function and suggest that increased astrocyte reactivity associated with early AD may contribute to cognitive decline and thus warranting further investigation.

More recently, sex differences in glial cell immune response have become increasingly recognized as a potential contributor to AD risk^5,7^. Increased astrocyte reactivity, as measured by plasma GFAP, has been reported in CU females at risk for future AD as compared with males^32^. Increased astrocyte reactivity has also been found in females in both community-based and large clinic cohorts that included a range of CU, MCI, and dementia participants^33–35^, as well as in a much larger population-based sample consisting of CU and MCI/dementia individuals^36^. Some studies have not reported sex-related differences in GFAP levels^37–39^, or stronger associations between Aβ and p-Tau in astrocyte-reactive males compared with females^40^. Importantly, few studies have examined whether sex modifies the association of GFAP with longitudinal cognition and AD biomarkers, such as plasma Aβ42/40, especially in individuals who are CU or without significant AD pathology^41^.

In the present study, we investigated sex differences in plasma GFAP and its longitudinal associations with plasma AD biomarkers and cognition in CU older adults from two cohorts: the community MERI program and the ADNI. Our primary aim was to determine whether sex modifies the association between GFAP and longitudinal global and domain-specific (e.g., episodic memory) cognitive decline. We conducted a hypothesis-driven examination of CU based on the prior biomarker studies that established that increased astrocytic reactivity may be in response to early AD pathology (Aβ+)^42^, including those who are Aβ-^18^, and may contribute to more rapid tau accumulation ^40^. We first examined biomarker associations providing the biological context for cognitive outcomes. We hypothesized that baseline plasma GFAP levels would be higher in females than in males and that higher GFAP would be associated with greater amyloid burden (reflected by lower plasma Aβ42/40) and increased tau pathology (reflected by elevated plasma p-Tau_231_), and that higher baseline GFAP and/or steeper longitudinal GFAP increases would predict greater declines in global cognition and domain-specific performance, more strongly in females.

## 2 METHODS

### 2.1 Population

#### 2.1.1 MERI Cohort

Participants were part of the Memory Education and Research Initiative (MERI), an ongoing, longitudinal observational cohort of community-dwelling individuals referred for memory evaluation. The MERI program is open to adults aged 18 years and older from Rockland County, New York, and surrounding areas (e.g., Westchester County, NY and Bergen County, NJ), but primarily targets older adults aged 50 and above. Participants are often referred by local physicians for assessment of cognitive impairment, while others are self-referred due to subjective cognitive concerns or a family history of AD or dementia. The MERI cohort includes more than 1,300 individuals. The present analysis focused on participants who provided blood samples during at least one MERI visit, and were classified as CU, using a broad cutoff of Mini-Mental State Examination (MMSE) ≥24^43^. Longitudinal analyses required at least one follow-up visit, the average number of follow-up visits was 3 (range 2-10). For certain analyses, participants with missing data were excluded as detailed below (Figure 1).

**Figure 1.**
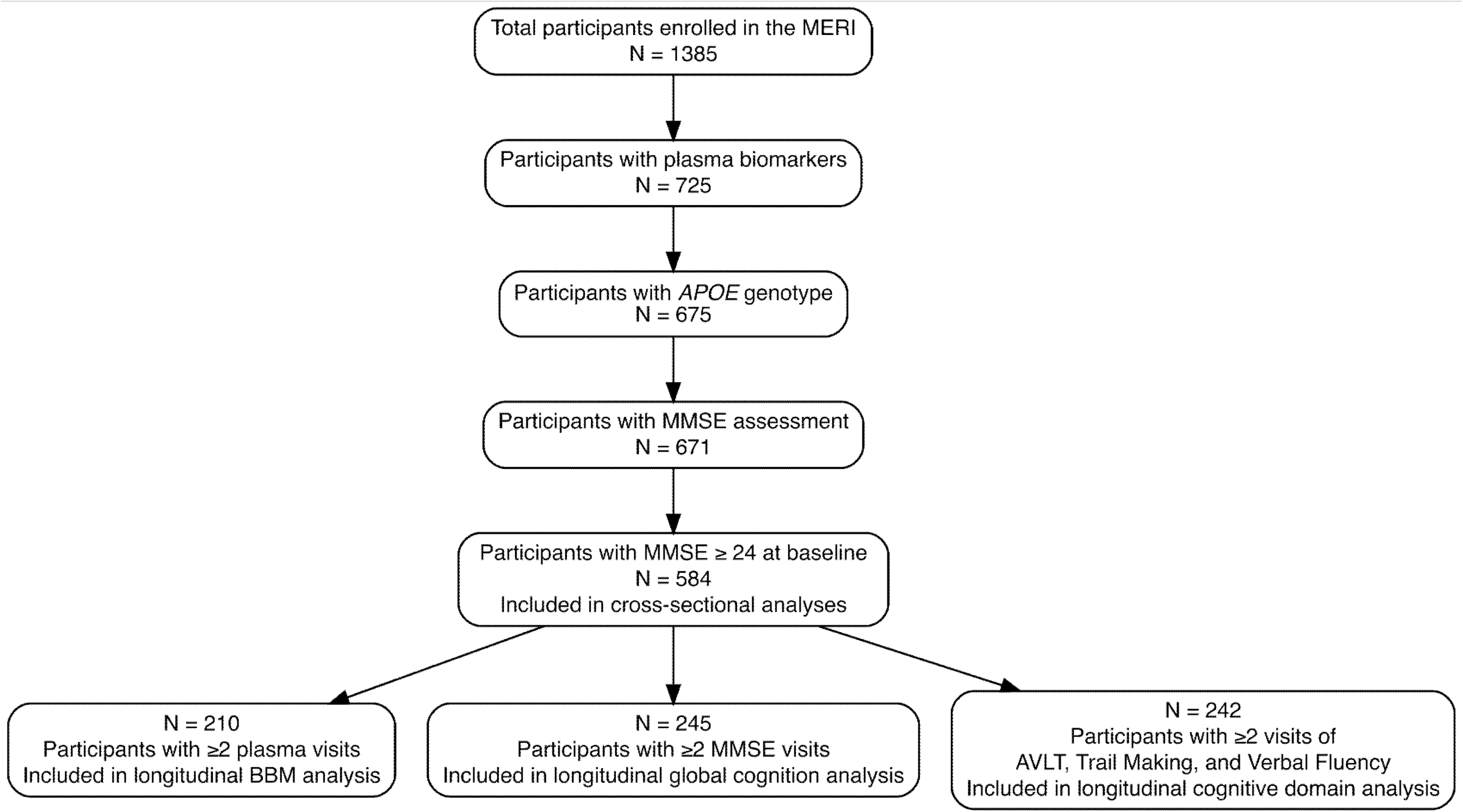
Participant flow diagram. The total number of MERI participants considered for this analysis is indicated in the top box. Each subsequent box describes the number of MERI participants, out of the total, which had plasma biomarkers, *APOE* genotype and MMSE ≥ 24. The number of MERI participants included in each of the cross-sectional analyses is described in the bottom boxes for the BBM analysis, global cognition analysis, and the cognitive domain analysis.

Most individuals enrolled in the MERI program were cognitively unimpaired (CU), healthy adults with stable medical conditions, as determined during the medical history.

#### 2.1.2 ADNI Cohort

The Alzheimer’s Disease Neuroimaging Initiative, a research-recruited cohort, was used for replication. Data used in the preparation of this article were obtained from the Alzheimer’s Disease Neuroimaging Initiative (ADNI) database (adni.loni.usc.edu). The ADNI was launched in 2003 as a public-private partnership, led by Principal Investigator Michael W. Weiner, MD. The primary goal of ADNI has been to test whether serial magnetic resonance imaging (MRI), positron emission tomography (PET), other biological markers, and clinical and neuropsychological assessment can be combined to measure the progression of mild cognitive impairment (MCI) and early Alzheimer’s disease (AD). For up-to-date information, see www.adni-info.org.

The ADNI recruited individuals between the ages of 55 to 90 years old across several waves [ADNI1, ADNI2, ADNI3, ADNI4]; the full inclusion/exclusion criteria can be found at ADNI website. The present analysis focused on ADNI participants who were from all waves who had *APOE* genotype, the biomarkers of interest, and were deemed CU based on the previously described criteria^44^. In brief, participants were included if the MMSE ≥ 24 and clinical dementia rating global score of 0 (Supplemental Figure 1). This ADNI cohort was included as a replication cohort.

### 2.2 Study Design

The MERI is a longitudinal study described in detail elsewhere^45^. In brief, participants complete the MERI program annually. Each visit consists of three components: a medical evaluation, a neuropsychological assessment, and a psychiatric evaluation. The medical evaluation includes a detailed history of past and current medical conditions and an optional blood draw. The neuropsychological assessment comprises 13 standardized tests covering several cognitive domains, including general intelligence and episodic verbal memory. For the present analysis, we selected tests sensitive to early and progressive changes along the AD spectrum^46^. The cognitive domains included in this analysis were episodic memory (Auditory Verbal Learning Test [AVLT] Total Learning and AVLT Delayed Recall), attention (Trail Making Test A time), executive function (Trail Making Test B time and Verbal Fluency), and semantic knowledge (Category Fluency). The psychiatric evaluation consists of a clinician-conducted interview and the assessment of psychiatric disorders, including depression and anxiety.

### 2.3 Blood Biomarkers

#### 2.3.1 MERI Blood Biomarkers

Non-fasting blood samples are collected in standard 10 mL EDTA tubes (BD Vacutainer) and processed as previously described^47^. In brief, blood was collected, kept at room temperature, and centrifuged within one hour of collection. The resulting plasma was transferred to pure polypropylene tubes and stored at -80 °F. The current study included MERI participants whose plasma samples had not been thawed since initial collection and storage. The first visit with an available blood-based biomarker (BBMs) sample, regardless of visit number, was designated as the baseline for subsequent AD biomarker analyses. Plasma concentrations of Aβ40, Aβ42, GFAP and NfL were measured using the Neuro-4-Plex-E Single molecule array (Simoa) assay (Quanterix). Plasma p-Tau231 concentration was measured using an in-house Simoa assay, as previously described^48^.

#### 2.3.2 ADNI Blood Biomarkers

Fasting blood samples were collected during the ADNI study visits, placed on dry ice, and were shipped for analysis on the day a sample was collected. Methodology is described on the ADNI website (http://www.adni-info.org/). Plasma concentrations of Aβ40, Aβ42, p-tau217, GFAP and NfL were measured using the Fujirebio Lumipulse G1200 and Quanterix HD-X^49^.

### 2.4 Statistical Analysis

All statistical analyses were performed in R (ver. 4.3.3). In the MERI cohort, GFAP, NfL, and p-Tau231 values were log10-transformed to follow a normal distribution, whereas Aβ values were not transformed. In the ADNI cohort, all BBMs were log10-transformed. Cognitive performance was assessed both globally (total MMSE score) and by cognitive domain (episodic memory, attention, executive function, and semantic knowledge). APOE genotype was categorized as E2 carriers (one or two E2 alleles), E3 homozygotes (two E3 alleles), or E4 copies (one or two E4 alleles). Baseline characteristics and group differences were compared using t-tests. For continuous variables, an ANOVA for multi-group comparisons (e.g., genotypes) and a t-test for two-group comparisons (e.g., sex) were performed. For categorical variables, Pearson’s chi-squared test was performed. Linear regression models (LMs) were applied to cross-sectional analyses at baseline while longitudinal analyses were performed using linear mixed models (LMMs; the *lmerTest* package in R).

Depending on the model, the primary exposure was defined either as a baseline measure (e.g., baseline GFAP) or as a time-varying longitudinal measure (e.g., longitudinal GFAP). LMMs included follow-up duration (time) to account for variability in across participants as well as baseline age as a covariate. Random intercepts in all LMMs account for within-subject correlations across repeated measures. All cross-sectional and longitudinal models were adjusted for age (or baseline age and time since baseline for LMMs), sex, education, APOE4 status, and race (non-Hispanic white vs other). To assess whether associations between GFAP and other BBMs or cognitive outcomes differed by sex, all models were additionally run within sex-stratified subgroups analyses followed by analyses with sex interaction terms. For longitudinal models with baseline GFAP as the exposure, the GFAP × time term tested whether the rate of change in the outcome varied according to baseline GFAP, whereas the GFAP × sex × time term formally tested whether this longitudinal association differed by sex. Lastly, analyses were performed to evaluate associations between within-individual rates of change (Δ per unit time) in GFAP and other BBMs, with models otherwise specified as above but adjusted for concurrent age and time between visits. Figures were generated using the DiagrammeR or ggplot2 package in R. Axis limits of figures for each biomarker or test were set to the same range for all figures to aid visualization; no data were excluded. All biomarkers were visually inspected for outliers and assessed normality. Mediation analysis provides us another way to investigate the sex effect on the association between GFAP and BBM or cognitive outcomes. Separate models were fit for each BBM and cognitive outcome using the framework sex → GFAP → outcome at baseline. The mediator model estimated the association between sex and GFAP, and the outcome model included both sex and GFAP controlling for age, race, education, and APOE4 status. Confidence intervals and p-values were estimated using 5,000 nonparametric bootstrap resamples.

## 3 RESULTS

### 3.1 Baseline Characteristics

#### 3.1.1 MERI Cohort

Our sample consisted of 584 cognitively unimpaired older adults enrolled in MERI. Baseline participant characteristics are presented in Table 1. Although females were more numerous than males, the two groups did not differ in demographic variables, including the proportion of *APOE4* carriers. We first examined sex differences in BBMs levels (Figure 2 a-f). Females showed significantly lower plasma concentrations of Aβ40 (p ≤ 0.001), Aβ42 (p = 0.012), and p-Tau231 (p = 0.002), and significantly higher plasma GFAP levels (p ≤ 0.001) compared with males. No significant sex differences were observed in baseline plasma Aβ42/40 ratio or NfL levels. Supplemental Table 1 presents performance on the MERI cognitive tests (MMSE and domain-specific assessments). At baseline, females performed better than males across all cognitive tests, except for the MMSE and the verbal fluency test.

**Figure 2.**
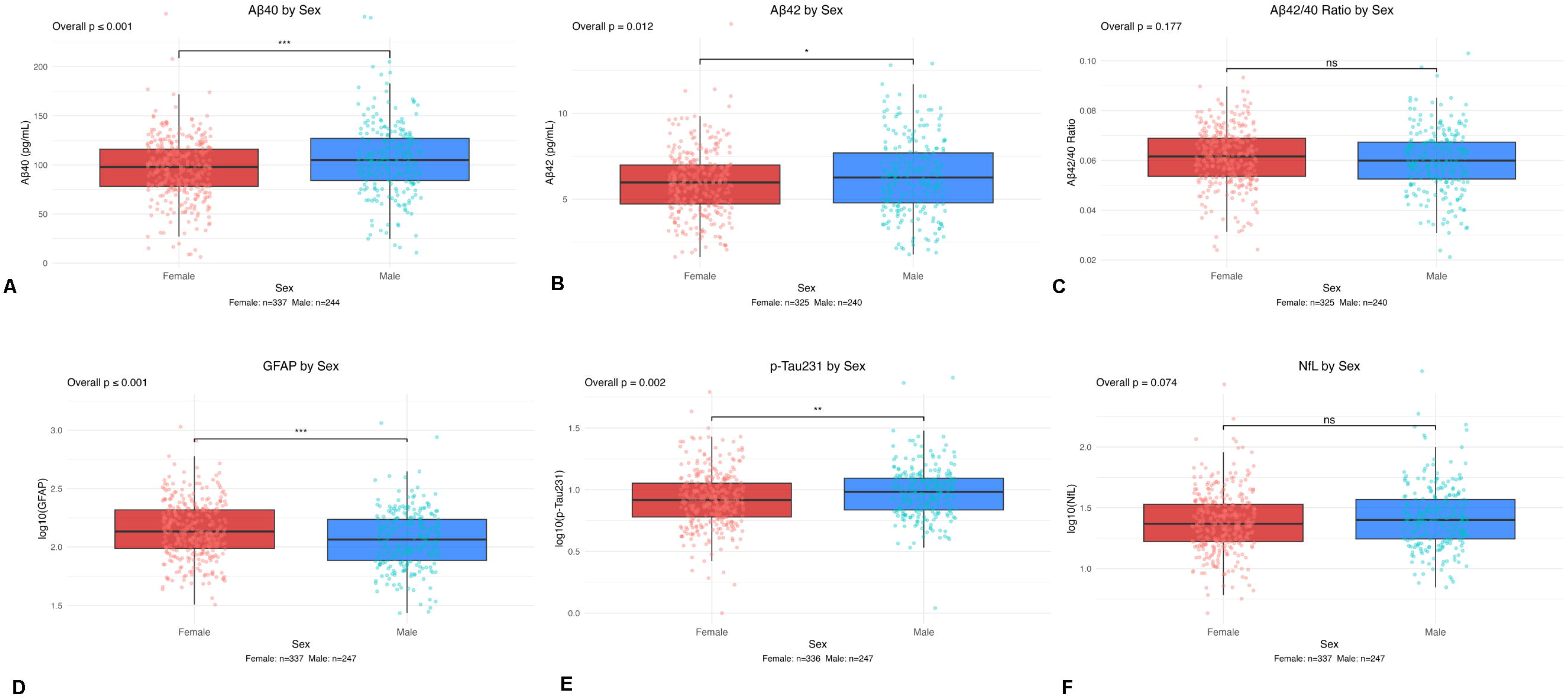
Boxplots of Blood-based Biomarkers (BBMs). Boxplots of Baseline MERI Cohort with individual points superimposed. Comparisons were plotted by sex (female: red, male: blue): Aβ40 (A), Aβ42 (B), Aβ42/40 (C), GFAP (D), p-Tau231 (E), and NfL (F). Significant associations are indicated with an asterisks (p<0.05*, p<0.01**, p<0.001***) and p > 0.05 indicated with ns.

**Table 1.**
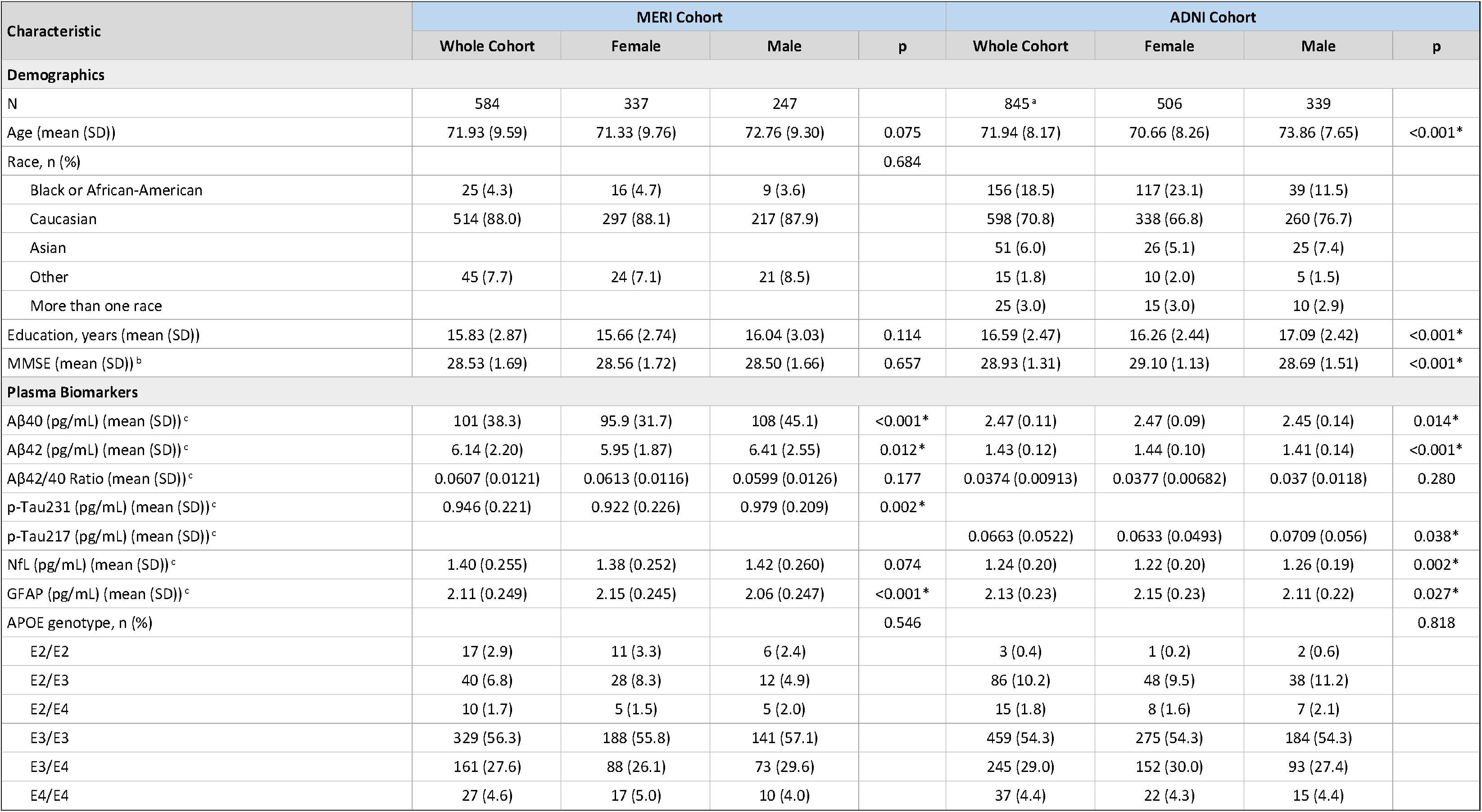
Cohort characteristics at baseline are presented for the whole sample and stratified by sex (female, male) for the MERI cohort and ADNI cohort. Group comparisons by sex were conducted using independent-samples t-tests. Statistical significance was set at p < 0.05 and is indicated by an asterisk (*).

#### 3.1.2 ADNI Cohort

Table 1 also reports for the ADNI cohort. Similar to the MERI cohort, females were more numerous than males. Males were significantly older than females, while females had a higher average level of education than males. The two groups did not differ in the proportion of *APOE4* carriers. We first examined sex differences in BBMs levels (Supplemental Figure 2 a-f). Females showed significantly higher plasma concentrations of Aβ40 (p = 0.014), Aβ42 p ≤ 0.001), p-Tau_217_ (p = 0.038), GFAP (p = 0.027), and NfL levels (p = 0.002) compared with males. No significant sex differences were observed in the baseline plasma Aβ42/40 ratio. Of note, while the direction of sex differences for Aβ40, Aβ42, and plasma p-Tau differed between the two cohorts (women lower in MERI, women higher in ADNI), the elevation of plasma GFAP in women and the absence of sex differences in the plasma Aβ42/40 ratio were consistent across both cohorts.

### 3.2 Associations Between Baseline GFAP and BBMS

#### 3.2.1 Baseline associations – MERI Cohort

Supplemental Table 2 reports LM analyses of the associations between baseline GFAP levels and baseline BBMs (Supplemental Figure 3). In the full MERI cohort, baseline GFAP was positively associated with Aβ40 (p < 0.001), Aβ42 (p = 0.002), p-Tau231 (p < 0.001), and NfL (p < 0.001), and negatively associated with the Aβ42/40 ratio (p < 0.001). In sex-stratified analyses, these associations were generally maintained; the Aβ42/40 ratio association was significant in females but not males, whereas the Aβ42 association was significant in males but not females (Supplemental Table 2). Note that GFAP × sex interaction results are in Supplemental Table 3.

#### 3.2.2 Baseline associations – ADNI Cohort

In the full ADNI cohort, baseline GFAP was positively associated with Aβ40 (p = 0.008), p-Tau217 (p < 0.001), and NfL (p < 0.001) (Supplemental Table 4). In sex-stratified analyses, the associations with p-Tau217 and NfL remained significant in both sexes, whereas the association with Aβ40 was significant in females but not males. Baseline GFAP was not significantly associated with Aβ42 or the Aβ42/40 ratio in either sex.

#### 3.2.3 Longitudinal outcomes – MERI Cohort

LMM analyses examined the associations between baseline GFAP levels and longitudinal BBM trajectories. Participants completed three to four follow-up visits on average. In models, no significant GFAP × sex × time or GFAP × sex interactions were observed for any BBM in MERI (Table 2 and Supplemental Table 5) indicating no statistical evidence that GFAP-related longitudinal BBM trajectories differed by sex . Therefore, these interaction terms are removed in the further analyses.

**Table 2.** Associations between baseline GFAP and longitudinal BBMs for the interaction between baseline GFAP and sex in a) the MERI cohort and b) the ADNI cohort. Output of linear mixed effects models. N represents unique timepoints; Participants (unique) represents the number of unique participants. Note that females are the reference group in the interaction. Each model was adjusted for age at baseline, time since baseline, race, education, and APOE4 status. Significant associations are indicated with asterisks (p<0.05*, p<0.01**, p<0.001***).

| Outcome (longitudinal) | N | Participants (unique) | Primary Exposure | Primary Exposure Beta (SE) | Primary Exposure p-value | Interaction Term | Interaction Beta (SE) | Interaction p-value | Significant Covariates |
|---|---|---|---|---|---|---|---|---|---|
| a) MERI Cohort |  |  |  |  |  |  |  |  |  |
| Aβ40 | 605 | 210 | GFAP (baseline) | 20.943 (10.729) | 0.052 | GFAP:Sex | 2.607 (14.316) | 0.856 | age at baseline, time since baseline |
| Aβ42 | 605 | 210 | GFAP (baseline) | -0.014 (0.710) | 0.984 | GFAP:Sex | 0.841 (0.948) | 0.376 | age at baseline, education, APOE4, time since baseline |
| Aβ42/40 | 605 | 210 | GFAP (baseline) | -0.012 (0.005) | 0.012* | GFAP:Sex | 0.008 (0.006) | 0.214 | APOE4 |
| p-Tau231 | 605 | 210 | GFAP (baseline) | 0.214 (0.066) | 0.001** | GFAP:Sex | -0.091 (0.089) | 0.307 | age at baseline, APOE4, time since baseline |
| NfL | 605 | 210 | GFAP (baseline) | 0.350 (0.076) | <0.001*** | GFAP:Sex | 0.014 (0.102) | 0.891 | age at baseline, race, time since baseline |
| b) ADNI Cohort |  |  |  |  |  |  |  |  |  |
| Aβ40 | 675 | 294 | GFAP (baseline) | 0.061 (0.029) | 0.035 | GFAP:Sex | -0.036 (0.043) | 0.401 | age at baseline, education, APOE4, time since baseline |
| Aβ42 | 675 | 294 | GFAP (baseline) | 0.023 (0.026) | 0.381 | GFAP:Sex | -0.033 (0.038) | 0.392 | age at baseline, education, APOE4, time since baseline |
| Aβ42/40 | 675 | 294 | GFAP (baseline) | -0.003 (0.002) | 0.170 | GFAP:Sex | 0.001 (0.003) | 0.832 | APOE4, time since baseline |
| p-Tau217 | 675 | 294 | GFAP (baseline) | 0.082 (0.018) | <0.001*** | GFAP:Sex | 0.023 (0.026) | 0.368 | age at baseline, race, APOE4, time since baseline |
| NfL | 548 | 294 | GFAP (baseline) | 0.279 (0.051) | <0.001*** | GFAP:Sex | 0.102 (0.075) | 0.175 | age at baseline, race, time since baseline |

In the full MERI cohort, higher baseline GFAP was associated with higher longitudinal levels of Aβ40 (p = 0.005), p-Tau_231_ (p < 0.001), and NfL (p < 0.001), and with lower longitudinal Aβ42/40 (p = 0.022) (Figure 3 and Table 4). In sex-stratified analyses, only NfL remained significantly associated with baseline GFAP in males, whereas in females significant associations were observed for Aβ40, Aβ42/40, p-Tau231, and NfL (Table 4). GFAP × time interaction was not significant for these outcomes in the full cohort (Supplemental Table 5). In sex-stratified analyses, the GFAP × time interaction was significant only for the Aβ42/40 ratio in males (β = 0.001, p = 0.042); no significant GFAP × time interactions were observed in females. Thus, although higher baseline GFAP tracked with an adverse BBM profile, there was limited evidence that baseline GFAP predicted the subsequent rate of BBM change in MERI. GFAP × age and GFAP × APOE ε4 terms are reported in Supplemental Tables 13 and 14.

**Figure 3.**
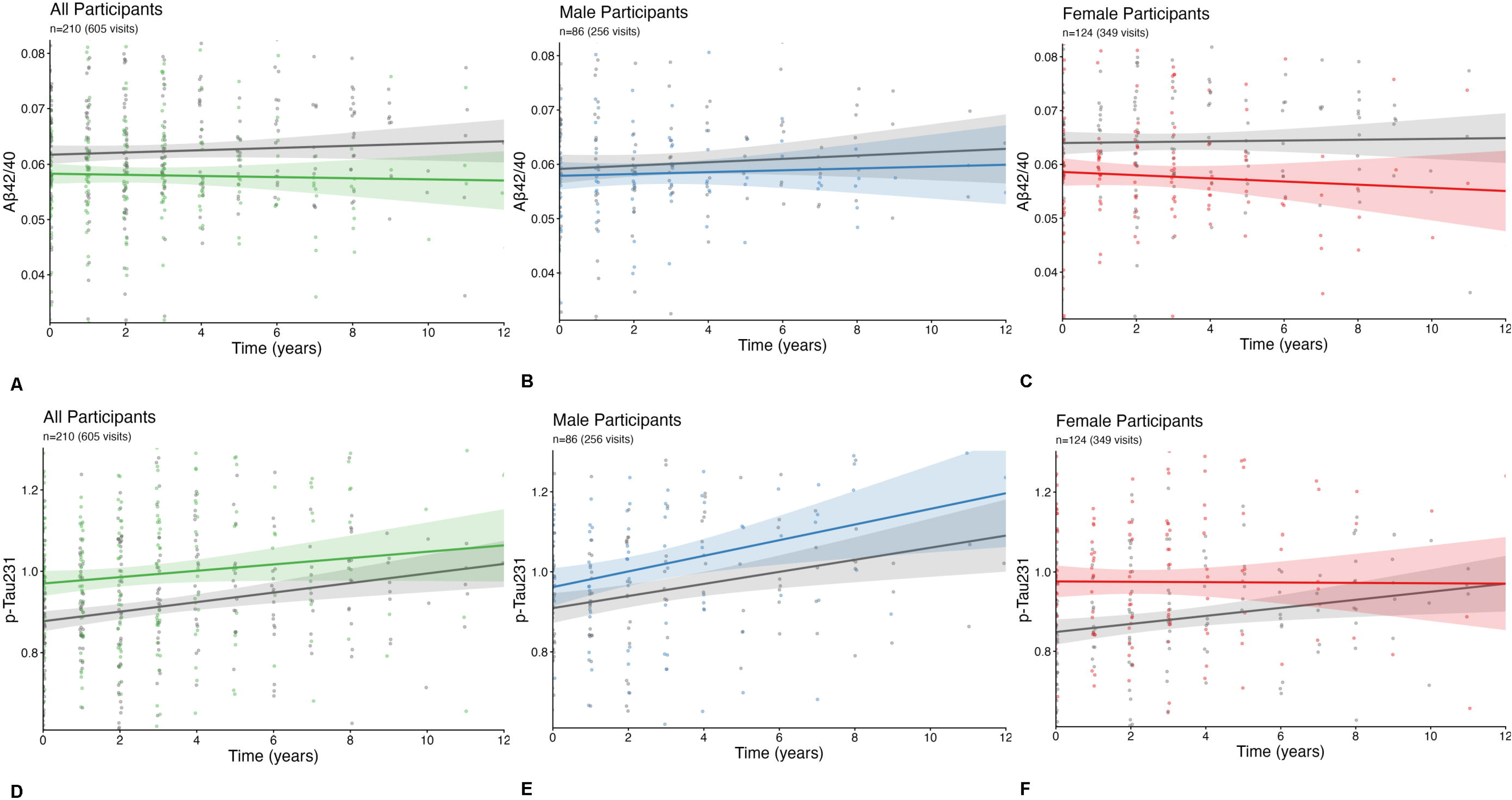
Longitudinal Aβ42/40 and p-Tau231 trajectories of participants in the lower vs. upper median GFAP at baseline for the MERI Cohort. Unadjusted linear trendlines depict longitudinal plasma obtained throughout follow up visits, comparing participants in the lower GFAP median (gray shading) versus upper GFAP median (colored shading) at baseline. Individual observations superimposed over trendlines. Panels show the full cohort (A,D), male participants only (B, E), and female participants only (C, F). Plots A-C are for Aβ42/40 ratio and D-F are for p-Tau231. Y-axis is plasma variable and X-axis is time (years). Plots were constructed from the subset of 210 participants with 2+ visits where plasma Aβ42/40/p-Tau231.

**Figure 4.**
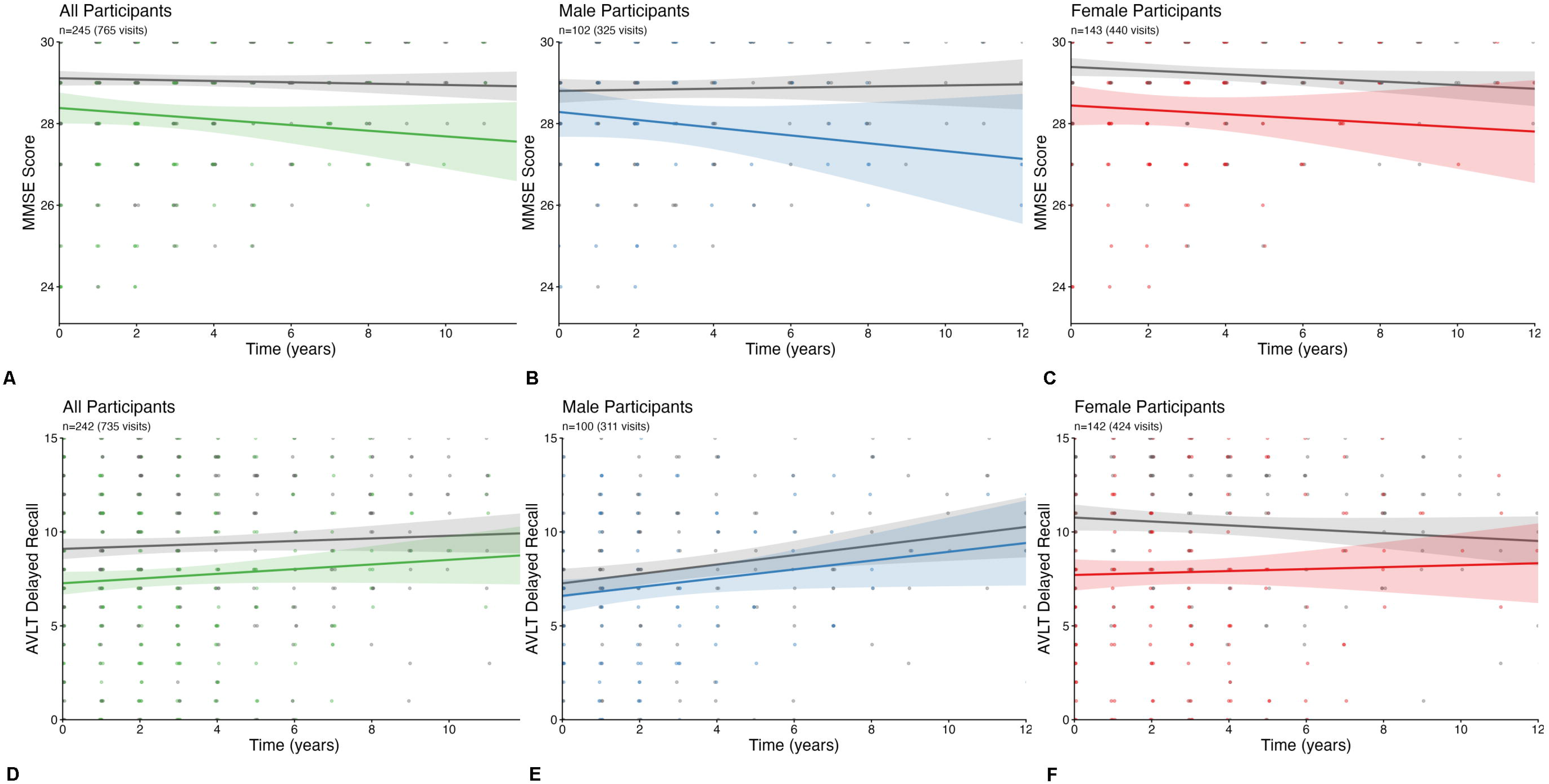
Longitudinal MMSE and AVLT score trajectories of female participants in the lower vs. upper median GFAP at baseline for the MERI Cohort. Unadjusted linear trendlines depict longitudinal MMSE scores across follow-up, comparing participants in the lower GFAP median (gray shading) versus upper GFAP median (colored shading) at baseline. Individual observations superimposed over trendlines. Panels show the full cohort (A, D), male participants only (B, E), female participants only (C, F). Plots A-C are for MMSE and D-F are for domain-specific cognition (AVLT Delayed Recall). Plots A-C are for MMSE and D-F are for domain-specific cognition (AVLT Delayed Recall). Plots were constructed from the subset of 245 participants with 2+ visits where MMSE or AVLT was completed. Y-axis is total score and X-axis is time (years).

#### 3.2.4 Longitudinal outcomes – ADNI Cohort

In ADNI cohort, participants completed two follow-up visits on average. No significant GFAP × sex interactions were observed for any BBM trajectories (Table 2). In the full cohort, baseline GFAP was associated with higher longitudinal levels of Aβ40, p-Tau217 (p < 0.001), and NfL (p < 0.001), while the GFAP × time interaction was significant only for p-Tau217 (β = 0.005, p = 0.015), indicating a steeper increase in p-Tau217 among participants with higher baseline GFAP (Supplemental Table 7b). In sex-stratified analyses, this GFAP × time association with p-Tau217 was significant in females (β = 0.007, p = 0.008) but not males (p = 0.924). However, the formal GFAP × sex × time interaction for p-Tau217 was not significant (p = 0.102), and no other BBM showed a significant three-way interaction (Supplemental Table 7). Baseline GFAP was not associated with a different longitudinal trajectory of the Aβ42/40 ratio. GFAP × age and GFAP × APOE ε4 terms are reported in Supplemental Tables 15 and 16.

### 3.3 Associations Between Baseline GFAP and Cognition

#### 3.3.1 Baseline associations – MERI Cohort

LMs examine the associations between GFAP levels and cognitive performance at baseline (Supplemental Figure 3). In the full MERI cohort (Supplemental Table 2), baseline GFAP was negatively associated with MMSE, AVLT Total Learning, AVLT Delayed Recall, verbal letter fluency, and verbal category fluency, and positively associated with Trail Making Test A and B completion times. In sex-stratified analyses, higher GFAP was associated with lower MMSE and AVLT Total Learning scores in males, whereas in females it was associated with lower MMSE, AVLT Total Learning, AVLT Delayed Recall, and category fluency scores (Supplemental Table 2). GFAP × sex interactions are reported in Supplemental Table 3.

#### 3.3.2 Baseline associations – ADNI Cohort

In ADNI, there were no significant associations between baseline GFAP and global cognition or domain-specific cognitive measures in the full cohort or in sex-stratified analyses (Supplemental Table 4). GFAP × sex interactions are reported in Supplemental Table 3.

#### 3.3.3 Longitudinal outcomes – MERI Cohort

We examined the associations between baseline GFAP levels and longitudinal changes in global cognition. In the full MERI cohort, higher baseline GFAP was associated with lower longitudinal MMSE scores (p = 0.001) (Table 3). The GFAP × sex × time interaction was not significant (p = 0.263). The GFAP × sex interaction for MMSE was not significant (Table 4). However , the GFAP × time interaction was significant for MMSE (β = -0.359, p < 0.001), indicating that higher baseline GFAP was associated with steeper MMSE decline over time (Supplemental Table 8b). This association was significant in both males (β = -0.534, p < 0.001) and females (β = -0.321, p = 0.003) (Supplemental Table 8).

**Table 3.** Associations between baseline GFAP and BBMs, MMSE, and cognitive performance in a) the full MERI cohort, b) males only, and c) females only. Output of linear mixed effects models. N and visit totals are reported once per outcome domain, as sample composition differs slightly by domain due to missing data. Significant associations are indicated with asterisks (p<0.05*, p<0.01**, p<0.001***).

| Outcome (longitudinal) | a) Full Cohort |  | b) Males Only |  | c) Females Only |  |
|---|---|---|---|---|---|---|
|  | Baseline GFAP, β (SE) | Baseline GFAP, p-value | Baseline GFAP, β (SE) | Baseline GFAP, p-value | Baseline GFAP, β (SE) | Baseline GFAP, p-value |
| **Blood-Based Biomarkers** | **N=210 participants across 605 total visits** |  | **N=86 participants across 256 total visits** |  | **N=124 participants across 349 total visits** |  |
| Aβ40 | 22.280 (7.819) | 0.005** <sup>a,f</sup> | 14.263 (12.703) | 0.265<sup>a</sup> | 30.102 (9.716) | 0.002** <sup>f</sup> |
| Aβ42 | 0.415 (0.520) | 0.425<sup>a,d,e,f</sup> | 0.764 (0.825) | 0.358<sup>—</sup> | -0.097 (0.686) | 0.887<sup>a,e,f</sup> |
| Aβ42/40 | -0.008 (0.003) | 0.022* <sup>e</sup> | -0.000 (0.005) | 0.997<sup>a,e</sup> | -0.016 (0.005) | 0.001*** <sup>e</sup> |
| p-Tau231 | 0.168 (0.049) | <0.001*** <sup>a,b,e,f</sup> | 0.105 (0.071) | 0.142<sup>a,f</sup> | 0.237 (0.069) | <0.001*** <sup>e,f</sup> |
| NfL | 0.357 (0.056) | <0.001*** <sup>a,c,f</sup> | 0.360 (0.087) | <0.001*** <sup>a,f</sup> | 0.336 (0.077) | <0.001*** <sup>a,f</sup> |
| **MMSE** | **N=245 participants across 765 total visits** |  | **N=102 participants across 325 total visits** |  | **N=143 participants across 440 total visits** |  |
| MMSE | -1.964 (0.566) | 0.001** <sup>a,b,c,d,f</sup> | -1.466 (0.828) | 0.080<sup>f</sup> | -2.623 (0.816) | 0.002** <sup>f</sup> |
| **Cognitive Testing** | **N=242 participants across 735 total visits** |  | **N=100 participants across 311 total visits** |  | **N=142 participants across 424 total visits** |  |
| AVLT, Total Learned | -9.279 (3.283) | 0.005** <sup>a,b,c,d,f</sup> | -5.112 (4.784) | 0.288<sup>a,f</sup> | -13.042 (4.667) | 0.006** <sup>a,f</sup> |
| AVLT, Delayed Recall | -3.879 (1.185) | <0.001*** <sup>a,b,f</sup> | -1.255 (1.609) | 0.437<sup>—</sup> | -6.559 (1.746) | <0.001*** <sup>f</sup> |
| Trail Making A (Time) | 4.178 (4.381) | 0.341<sup>a,b,c,f</sup> | 8.019 (8.176) | 0.330<sup>f</sup> | 1.883 (4.873) | 0.700<sup>a,c,e,f</sup> |
| Trail Making B (Time) | 21.801 (16.612) | 0.191<sup>a,b,c,d,f</sup> | 42.505 (30.784) | 0.171<sup>a,f</sup> | 6.853 (17.747) | 0.700<sup>a,c,d,f</sup> |
| Trail Making (B-A) | 17.052 (13.534) | 0.209<sup>a,b,c,d,f</sup> | 33.837 (24.579) | 0.173<sup>a,f</sup> | 4.880 (14.769) | 0.742<sup>a,c,d,f</sup> |
| Trail Making (B/A) | 0.135 (0.285) | 0.636<sup>a,c,d,f</sup> | 0.300 (0.449) | 0.506<sup>a,f</sup> | -0.044 (0.369) | 0.906<sup>c,d,f</sup> |
| Verbal Fluency, Animals (Total) | -2.103 (1.488) | 0.159<sup>a,b,c,f</sup> | 0.314 (2.219) | 0.888<sup>a,c,f</sup> | -4.202 (2.045) | 0.042* <sup>a,c,f</sup> |
| Verbal Fluency, F+A+S (Total) | -2.197 (3.933) | 0.577<sup>b,c,d,f</sup> | 1.724 (5.810) | 0.767<sup>—</sup> | -5.407 (5.409) | 0.319<sup>c,d,f</sup> |

**Table 4.**
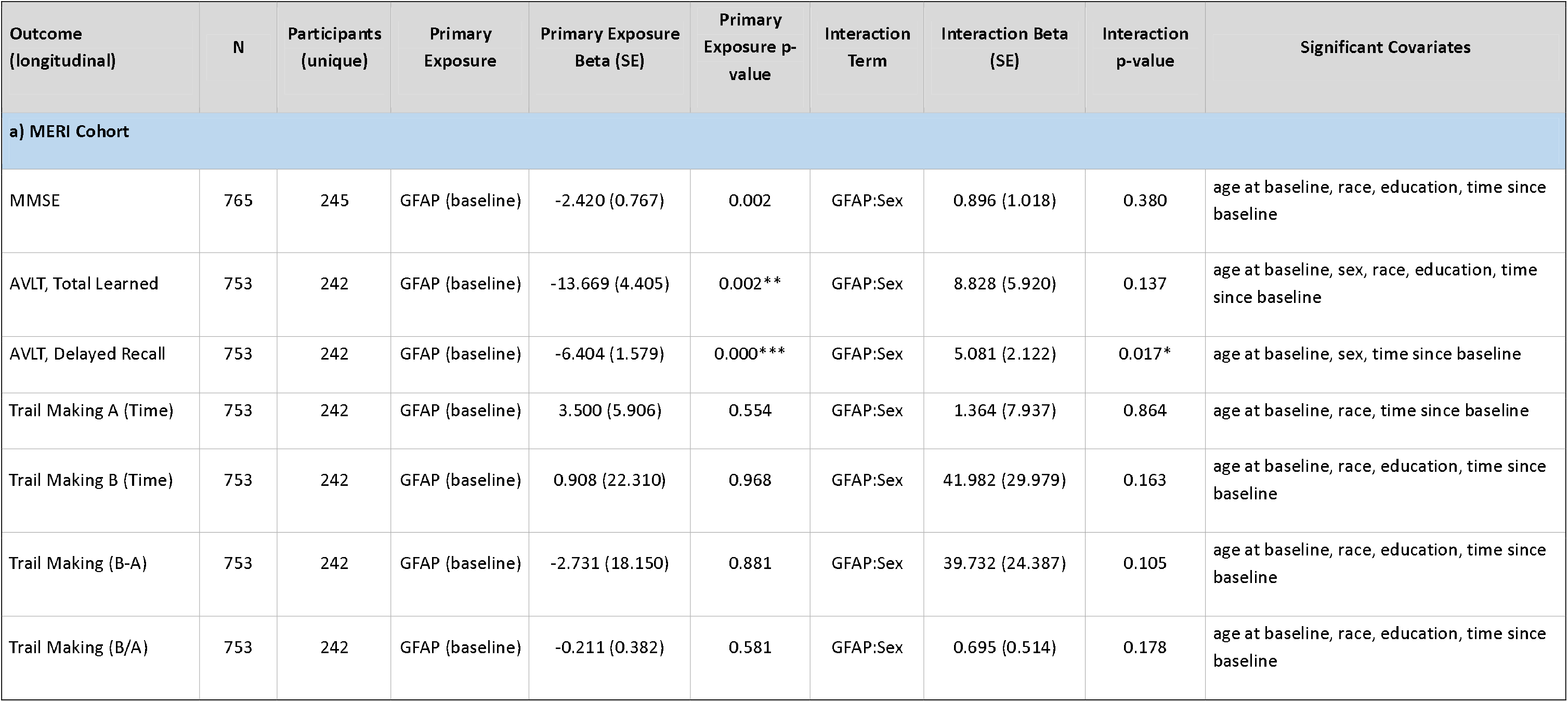
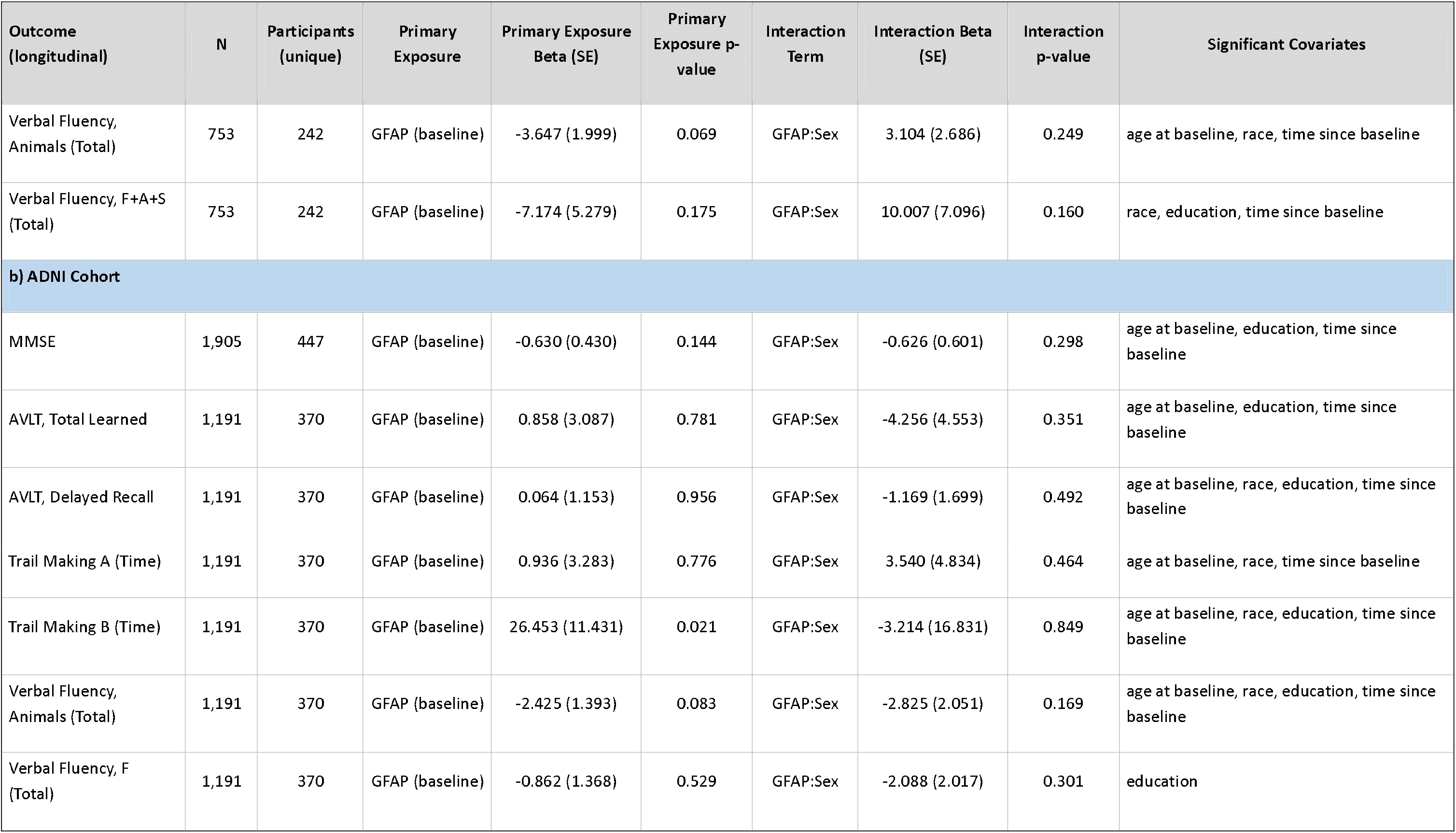
Associations between baseline GFAP and longitudinal global and domain-specific cognitive performance for the interaction between baseline GFAP and sex in a) the MERI cohort and b) the ADNI cohort. Output of linear mixed effects models. N represents unique timepoints; Participants (unique) represents the number of unique participants. Note that females are the reference group in the interaction. Each model was adjusted for age at baseline, time since baseline, race, education, and APOE4 status. Significant associations are indicated with asterisks (p<0.05*, p<0.01**, p<0.001***).

| Outcome (longitudinal) | N | Participants (unique) | Primary Exposure | Primary Exposure Beta (SE) | Primary Exposure p-value | Interaction Term | Interaction Beta (SE) | Interaction p-value | Significant Covariates |
|---|---|---|---|---|---|---|---|---|---|
| **a) MERI Cohort** |  |  |  |  |  |  |  |  |  |
| MMSE | 765 | 245 | GFAP (baseline) | -2.420 (0.767) | 0.002 | GFAP:Sex | 0.896 (1.018) | 0.380 | age at baseline, race, education, time since baseline |
| AVLT, Total Learned | 753 | 242 | GFAP (baseline) | -13.669 (4.405) | 0.002** | GFAP:Sex | 8.828 (5.920) | 0.137 | age at baseline, sex, race, education, time since baseline |
| AVLT, Delayed Recall | 753 | 242 | GFAP (baseline) | -6.404 (1.579) | 0.000*** | GFAP:Sex | 5.081 (2.122) | 0.017* | age at baseline, sex, time since baseline |
| Trail Making A (Time) | 753 | 242 | GFAP (baseline) | 3.500 (5.906) | 0.554 | GFAP:Sex | 1.364 (7.937) | 0.864 | age at baseline, race, time since baseline |
| Trail Making B (Time) | 753 | 242 | GFAP (baseline) | 0.908 (22.310) | 0.968 | GFAP:Sex | 41.982 (29.979) | 0.163 | age at baseline, race, education, time since baseline |
| Trail Making (B-A) | 753 | 242 | GFAP (baseline) | -2.731 (18.150) | 0.881 | GFAP:Sex | 39.732 (24.387) | 0.105 | age at baseline, race, education, time since baseline |
| Trail Making (B/A) | 753 | 242 | GFAP (baseline) | -0.211 (0.382) | 0.581 | GFAP:Sex | 0.695 (0.514) | 0.178 | age at baseline, race, education, time since baseline |
| Outcome (longitudinal) | N | Participants (unique) | Primary Exposure | Primary Exposure Beta (SE) | Primary Exposure p-value | Interaction Term | Interaction Beta (SE) | Interaction p-value | Significant Covariates |
| Verbal Fluency, Animals (Total) | 753 | 242 | GFAP (baseline) | -3.647 (1.999) | 0.069 | GFAP:Sex | 3.104 (2.686) | 0.249 | age at baseline, race, time since baseline |
| Verbal Fluency, F+A+S (Total) | 753 | 242 | GFAP (baseline) | -7.174 (5.279) | 0.175 | GFAP:Sex | 10.007 (7.096) | 0.160 | race, education, time since baseline |
| **b) ADNI Cohort** |  |  |  |  |  |  |  |  |  |
| MMSE | 1,905 | 447 | GFAP (baseline) | -0.630 (0.430) | 0.144 | GFAP:Sex | -0.626 (0.601) | 0.298 | age at baseline, education, time since baseline |
| AVLT, Total Learned | 1,191 | 370 | GFAP (baseline) | 0.858 (3.087) | 0.781 | GFAP:Sex | -4.256 (4.553) | 0.351 | age at baseline, education, time since baseline |
| AVLT, Delayed Recall | 1,191 | 370 | GFAP (baseline) | 0.064 (1.153) | 0.956 | GFAP:Sex | -1.169 (1.699) | 0.492 | age at baseline, race, education, time since baseline |
| Trail Making A (Time) | 1,191 | 370 | GFAP (baseline) | 0.936 (3.283) | 0.776 | GFAP:Sex | 3.540 (4.834) | 0.464 | age at baseline, race, time since baseline |
| Trail Making B (Time) | 1,191 | 370 | GFAP (baseline) | 26.453 (11.431) | 0.021 | GFAP:Sex | -3.214 (16.831) | 0.849 | age at baseline, race, education, time since baseline |
| Verbal Fluency, Animals (Total) | 1,191 | 370 | GFAP (baseline) | -2.425 (1.393) | 0.083 | GFAP:Sex | -2.825 (2.051) | 0.169 | age at baseline, race, education, time since baseline |
| Verbal Fluency, F (Total) | 1,191 | 370 | GFAP (baseline) | -0.862 (1.368) | 0.529 | GFAP:Sex | -2.088 (2.017) | 0.301 | education |

For domain-specific cognition, higher baseline GFAP was associated with poorer longitudinal AVLT Total Learning (p = 0.005) and AVLT Delayed Recall (p ≤ 0.001) performance in the full MERI cohort (Table 3). A significant GFAP × sex interaction was observed for AVLT Delayed Recall (p = 0.017), whereas the interaction for AVLT Total Learning was not significant (p = 0.137) (Table 4). In models testing change over time, higher baseline GFAP was associated with steeper longitudinal decline in AVLT Total Learning (GFAP × time: p = 0.021) and AVLT Delayed Recall (p < 0.001), and with worsening Trail Making Test B performance (p = 0.007) in the full cohort (Supplemental Table 8d). In sex-stratified analyses, the GFAP × time association with AVLT Delayed Recall was significant in both males (p = 0.035) and females (p < 0.001), whereas the association with AVLT Total Learning was significant in females (p = 0.007) but not males (p = 0.692). Worsening Trail Making Test B performance was also associated with baseline GFAP in both males (p = 0.033) and females (p = 0.030). We note that the GFAP × sex × time interactions for domain-specific cognitive outcomes were not significant (Supplemental Table 8c).

#### 3.3.4 Longitudinal outcomes – ADNI Cohort

In ADNI, no significant GFAP × sex interactions were observed for MMSE or any domain-specific cognitive outcome (Table 4). For MMSE, the GFAP × time interaction was significant in the full cohort (β = -0.190, p = 0.001), indicating steeper MMSE decline with higher baseline GFAP (Supplemental Table 10b). The association was significant in males (β = -0.212, p = 0.008) but not females (p = 0.266); the GFAP × sex × time interaction was not significant (p = 0.375) (Supplemental Table 10). For domain-specific outcomes, higher baseline GFAP was associated with worsening Trail Making Test B performance and reduced verbal category fluency over time in the full cohort (Supplemental Table 10d). Although some associations differed in significance in sex-stratified models, GFAP × sex × time interactions were significant for Trail Making Test A (p = 0.015) and verbal category fluency (p = 0.025), but not for the other domain-specific outcomes (Supplemental Table 10c). GFAP × age and GFAP × APOE ε4 analyses are reported in Supplemental Tables 15 and 16.

### 3.4 Longitudinal GFAP and Longitudinal BBM and Cognition analyses – MERI Cohort

Supplemental Table 11 reports the MERI analyses using longitudinal GFAP as a time-varying exposure: In the full cohort, higher longitudinal GFAP was associated with higher Aβ40, Aβ42, p-Tau231, and NfL levels (all p < 0.001) and with a lower Aβ42/40 ratio (p = 0.001). These BBM associations were broadly similar in sex-stratified analyses, although the Aβ42/40 association was significant in females (p = 0.003) but not males. Higher longitudinal GFAP was also associated with lower MMSE scores in the full cohort (p = 0.024) and in males (p = 0.027), but not females. For domain-specific cognition, higher longitudinal GFAP was associated with poorer AVLT Total Learning (p < 0.001), AVLT Delayed Recall (p = 0.003), and category fluency (p = 0.013) in the full cohort. In males, associations were also observed with Trail Making Test B and B-A performance, whereas in females significant associations were observed for AVLT Total Learning and category fluency.

### 3.5 Rates of Change in GFAP and Longitudinal BBM analyses

#### 3.5.1 MERI Cohort

Supplemental Table 12 reports the results of analyses on within-individual rates of change in GFAP and BBMs in MERI. Faster GFAP increase was associated with faster increases in Aβ40, Aβ42, p-Tau231, and NfL and with faster decreases in the Aβ42/40 ratio in the full cohort (all p ≤ 0.012). In males, associations persisted for Aβ40, Aβ42, Aβ42/40, and NfL, but not p-Tau231. In females, associations remained significant for Aβ40, p-Tau_231_, and NfL, whereas Aβ42 and Aβ42/40 were not significantly associated (both p < 0.10).

#### 3.5.2 ADNI Cohort

Faster GFAP increase was associated with faster increases in Aβ40, Aβ42, Aβ42/40, p-Tau217, and NfL in the full cohort (p < 0.001). In males, associations remained significant for Aβ42, Aβ42/40, p-Tau217, and NfL, but not Aβ40. In females, associations were significant for all BBMs (Supplemental Table 12).

### 3.6 Mediation Analyses

#### 3.6.1 MERI Cohort

Mediation analyses at baseline identified significant associations through GFAP for several BBMs and cognitive outcomes in MERI (Supplemental Table 17). Significant mediation effects were observed for p-Tau231, Aβ40, Aβ42, Aβ42/40, NfL, MMSE, both AVLT measures, Trail Making Test A and B, and both verbal fluency measures.

#### 3.6.2 ADNI Cohort

In ADNI, significant associations through GFAP were observed for p-tau217, Aβ40, Aβ42/40, and NfL, whereas no significant effects were observed for MMSE or domain-specific cognitive measures (Supplemental Table 17). Note that these analyses were considered indirect pathway.

## 4 DISCUSSION

We examined associations between plasma GFAP and other BBMs, and cognition in CU older adults from MERI and replicated the analyses in ADNI. Females had higher plasma GFAP in both cohorts, whereas sex differences in other BBMs were cohort-dependent: females had lower Aβ40, Aβ42, and p-tau231 in MERI but higher Aβ40, Aβ42, p-tau217, and NfL in ADNI. The Aβ42/40 ratio did not differ by sex in either cohort. In MERI, higher baseline GFAP was associated with lower Aβ42/40 and higher p-Tau231 levels, consistent with greater AD-related pathology^50^, with no significant sex×GFAP interactions for BBM outcomes. Baseline GFAP was associated with steeper decline in global cognition and episodic memory, with no significant GFAP × sex × time interactions. Thus, formal longitudinal interaction models provided limited evidence that these biomarker trajectories differed by sex, which may be due to variability in follow-up durations among participants. In ADNI, the inverse association between GFAP and Aβ42/40 and the association with longitudinal p-Tau217 increases were again evident in females, again with no significant sex×GFAP interactions for BBM. Baseline GFAP was associated with steeper global cognitive decline and with worsening executive and semantic measures; only selected domain-specific outcomes showed significant three-way interactions. We found that the consistent sex difference was higher GFAP in females, whereas sex-specific longitudinal associations were less consistent and require replication.

Previous studies show that plasma GFAP increases across the AD continuum, discriminates preclinical AD from MCI and dementia, predicts cognitive and functional decline in Aβ+individuals, and may rise years before clinical conversion^51,52^. Postmortem studies similarly demonstrate that higher GFAP levels are associated with greater AD pathology^53^. Our findings extend this work by showing that GFAP is related to longitudinal AD-associated plasma biomarker profile changes in CU individuals. However, the formal interaction analyses indicate that these longitudinal biomarker associations should not generally be interpreted as female-specific. We also observed longitudinal cognitive change, in women, which may signal increased vulnerability to future cognitive decline. Greater clinical progression was not observed and would be difficult to establish given the relatively short follow-up of approximately 3-4 years. Nonetheless, elevated GFAP is not specific to AD. Chronic inflammatory conditions^55,56^, major depressive disorder^57^, bipolar depression^58^, progressive multiple sclerosis^59^, severe infections^60^, and other systemic or neurological conditions may increase GFAP. Although MERI and ADNI participants were generally medically stable, comorbidities were not exclusionary and may have influenced GFAP levels.

While elevations in plasma GFAP have been consistently reported across the AD spectrum and in CU individuals^36,61^, only recently have studies begun to examine the influence of sex. Across the AD spectrum, plasma GFAP levels are elevated in women^22,32–35,37,62^, with only a few studies that have not observed sex differences in GFAP^42,63^. Women exhibit faster increases in GFAP over time, but this may depend on age^32^ as well as ε4 carrier status^34^. Other reports suggest sex differences in associations with GFAP that are related to tau accumulation and vary by disease stage^36,64^. CSF GFAP may also show different sex associations from plasma GFAP^65^. This heterogeneity likely reflects differences in cohort composition, including age range, Aβ positivity, cognitive status, and assay platform. MERI was enriched for individuals with cognitive concerns but classified as CU, whereas ADNI CU participants were enrolled through standardized research criteria. Against these differences, the replication of higher GFAP in women and of no sex difference in Aβ42/40 supports the robustness of these signals.

Our study extends this literature in several ways. First, our primary analyses evaluated a strictly CU cohort recruited through physician- and self-referral for cognitive concerns, thereby addressing an earlier and less clinically confounded stage of the AD continuum. Second, we examined baseline and longitudinal GFAP in relation to both BBM and cognitive trajectories within the same cohort and tested these findings in ADNI. Third, we complemented sex-stratified analyses with interaction models. This distinction facilitates a more intuitive identification of potentially meaningful effect modifications by sex, independent of the statistical significance of GFAP × sex interactions.

The absence of sex differences in plasma Aβ42/40 in both cohorts suggests that greater amyloid burden does not fully explain higher GFAP in women^50,66^. Sex differences in plasma tau were inconsistent, with higher p-Tau231 in men in MERI and higher p-Tau217 in women in ADNI, whereas larger community data reported no sex differences in p-Tau217^67^. These discrepancies indicate that sex effects on plasma tau in CU populations remain unsettled. Prior work suggests that astrocyte reactivity may influence the relationship between amyloid and tau, and that diagnostic performance improves when GFAP is combined with p-Tau217^40,68^. In our longitudinal ADNI analyses, higher baseline GFAP was associated with a steeper increase in p-Tau217As our study did not include PET imaging or derive sex-specific cutoffs^65^, conclusions about sex-specific-specific AD pathology remain speculative.

Higher GFAP was also associated with longitudinal cognitive decline. In MERI, higher baseline GFAP was associated with steeper decline in MMSE and episodic memory, with significant GFAP × time effects in both sexes for MMSE and AVLT Delayed Recall and no significant GFAP × sex × time interactions. In ADNI, higher baseline GFAP was associated with steeper MMSE decline and with worsening executive and semantic measures. Although some sex-stratified associations differed, formal three-way interactions were significant only for selected domain-specific outcomes, including Trail Making Test A and category fluency. Women had better baseline cognitive performance in both cohorts, consistent with prior work in older adults^69^ and CU ADNI participants^70^. Prior studies have linked higher GFAP to poorer global cognition^18,71–73^, episodic memory^26,73^, executive function^29,30^ and language abilities^28,30^. Therefore, the different domains implicated in MERI and ADNI likely reflect cohort characteristics, baseline performance, follow-up duration, and test sensitivity rather than divergent underlying biology.

Sex-specific glial and endocrine mechanisms may contribute to these findings^7^. Biological sex influences neuroinflammatory dynamics, and age-related endocrine changes affect astrocyte function^34,74^. Estrogen modulates neuroimmune response^75^, mitochondrial function^76^, aromatase-dependent microglial activity^77^, and neurotrophic signaling relevant to hippocampal activity^78^. Evidence on hormone replacement therapy (HRT) during menopause is mixed^74^, with dementia risk depending on formulation and timing^79–81^. Postmenopausal endocrine shifts may also alter microglial activation and astrocyte states^82^. Because most women in our cohorts were postmenopausal, GFAP elevations may partly reflect these mechanisms^83^. Moreover, these factors raise the possibility that increased astrocytic reactivity may contribute to AD risk in women through mechanisms involving amyloid production or impaired clearance^12^. However, menopause status, estrogen levels, and HRT use were not modeled, so this interpretation remains speculative.

The observation of higher GFAP in women despite no higher Aβ42/40 burden is clinically relevant. GFAP may help distinguish CU individuals with subjective complaints from Aβ– MCI^84^ and MERI was largely Aβ– based on plasma p-Tau231 cutoffs^85^. Plasma GFAP may be more closely linked to brain amyloid pathology than CSF GFAP, whereas CSF GFAP may reflect broader neuroglial injury^22,86^. Thus, plasma and CSF GFAP should not be treated as interchangeable^87^. Additionally, astrocyte activation is not always a response to primary elevations in brain Aβ; thus, may reflect a response to blood–brain barrier (BBB) dysfunction or systemic inflammation^84^. Recent work also cautions that plasma GFAP cannot be interpreted as a pure measure of brain astrocyte reactivity because BBB dysfunction and non-astrocytic sources may contribute^86,87^. Future studies using paired plasma and CSF sampling, PET, inflammatory markers, BBB measures, and estrogen-related measures are needed to clarify whether GFAP in CU women reflects early AD processes, systemic inflammation, BBB dysfunction, or overlapping mechanisms.

### 4.1 Limitations

Several limitations should be noted. First, MERI and ADNI were composed largely of highly educated, non-Hispanic White individuals, which limits generalizability. Second, women outnumbered men, although groups were broadly comparable and models adjusted for demographic variables and APOE. Previous studies suggest that plasma GFAP may not be strongly influenced by *APOE-*ε*4* genotype^88^, but *APOE* was included as a covariate. Third, GFAP increases with age, and age-related imbalance could influence results^12,89^. However, MERI did not show a sex difference in the proportion aged ≥65 years (p=0.36), and analyses were adjusted for age. Fourth, direct measures of AD pathology were unavailable in MERI. Plasma AD biomarkers correlate with amyloid and tau pathology but remain indirect measures^48,90^. Fifth, factors explaining GFAP variance including sex, only account for a small portion of variance in GFAP^91^. Additional factors like body mass index, kidney function, other comorbidities, lifestyle^91^, sleep quality and duration^92,93^, and air pollution^94^ were not fully modeled. Finally, MERI participants were classified as CU but were enriched for subjective complaints or family history of dementia through physician- and self-referral recruitment. Findings should be interpreted in the context of this ascertainment and validated in more diverse population-based cohorts.

We conjecture that inconsistent findings regarding interaction terms may be attributable to participant attrition across cohorts. Furthermore, variability in follow-up durations among participants complicates the construction of statistical models capable of yielding accurate inferences regarding interaction effects. Given the inherent limitations of large observational cohorts, stratified analyses by sex may provide a more robust approach to examining sex-specific effects. We were also unable to perform an ADNI CSF GFAP replication analysis because there was insufficient overlap between participants with CSF GFAP measurements and those with the longitudinal plasma data required for the present analyses. We did not perform analyses stratified by plasma Aβ42/40 positivity because the number of putatively positive MERI participants was small and a directly comparable validated positivity threshold was not available across the assays used in the two cohorts. Finally, comorbidity-stratified analyses were not performed because medical-history variables were not sufficiently harmonized across MERI and ADNI to define robust vascular or other comorbidity subgroups.

## 5 CONCLUSIONS

Across two independent cohorts, the most reproducible sex-related finding was higher plasma GFAP in women. GFAP was also associated with AD-related BBMs and subsequent cognitive decline, with sex-stratified analyses in both cohorts suggested female-predominant associations in both MERI and ADNI. Based on prior biomarker and mechanistic studies, elevated plasma GFAP may reflect astrocytic reactivity associated with early AD-related biological processes^95^. These associations are likely influenced by sex-related immune and hormonal factors^74^ may contribute to differences in astrocytic responses. This interpretation is compatible with prior work showing that astrocyte reactivity can influence the relationship between amyloid and tau pathology^40,96^. However, the present observational findings do not establish the temporal ordering or causal direction of these processes. Moreover, formal interaction analyses did not support a consistently female-specific effect of GFAP on longitudinal BBM or cognitive trajectories, noting that these inconsistent findings may be related to cohort differences and participant attrition across cohorts. Future studies incorporating direct measures of brain AD pathology, larger samples with equal number of follow-up visits powered for sex interactions, and relevant inflammatory and endocrine measures are needed to determine whether astrocytic reactivity contributes differently to preclinical processes, in women and men.

## ACKNOWLEDGEMENTS

We would like to acknowledge all the ADNI, the ADNI study sites and the ADNI participants for the use of the data analyzed as part of the replication analyses. HZ is a Wallenberg Scholar and a Distinguished Professor at the Swedish Research Council supported by grants from the Swedish Research Council (#2023-00356, #2022-01018 and #2019-02397), the European Union’s Horizon Europe research and innovation programme under grant agreement No 101053962, Swedish State Support for Clinical Research (#ALFGBG-71320), the Alzheimer Drug Discovery Foundation (ADDF), USA (#201809-2016862), the AD Strategic Fund and the Alzheimer’s Association (#ADSF-21-831376-C, #ADSF-21-831381-C, #ADSF-21-831377-C, and #ADSF-24-1284328-C), the European Partnership on Metrology, co-financed from the European Union’s Horizon Europe Research and Innovation Programme and by the Participating States (NEuroBioStand, #22HLT07), the Bluefield Project, Cure Alzheimer’s Fund, the Olav Thon Foundation, the Erling-Persson Family Foundation, Familjen Rönströms Stiftelse, Familjen Beiglers Stiftelse, Stiftelsen för Gamla Tjänarinnor, Hjärnfonden, Sweden (#FO2022-0270), the European Union’s Horizon 2020 research and innovation programme under the Marie Skłodowska-Curie grant agreement No 860197 (MIRIADE), the European Union Joint Programme – Neurodegenerative Disease Research (JPND2021-00694), the National Institute for Health and Care Research University College London Hospitals Biomedical Research Centre, the UK Dementia Research Institute at UCL (UKDRI-1003), and an anonymous donor.

## Declaration of generative AI use

During the preparation of this work the author(s) used ChatGPT 5.0 in order to check English grammar/style and improve readability. After using this tool/service, the author(s) reviewed and edited the content as needed and take(s) full responsibility for the content of the published article. During the revision of this paper, Anthropic (Claude Sonnet 5) was used to reformat and condense the number of tables for readability. After using this tool/service, the author(s) reviewed and edited for accuracy.

## Data Sharing and Code Availability

The MERI data associated with this publication is not publicly available. The original study protocol did not include a data sharing provision; therefore, participants did not agree for their data to be shared publicly. The code generated for this analysis is available upon request. The ADNI data associated with this publication is publicly available to approved researchers (https://adni.loni.usc.edu/data-samples/adni-data/).

## FUDNING SOURCES

This MERI program has been partially supported by the Office of Mental Health of Rockland County, NY awarded to NP.

Data collection and sharing for the Alzheimer’s Disease Neuroimaging Initiative (ADNI) is funded by the National Institute on Aging (National Institutes of Health Grant U19 AG024904). The grantee organization is the Northern California Institute for Research and Education.

In the past, ADNI has also received funding from the National Institute of Biomedical Imaging and Bioengineering, the Canadian Institutes of Health Research, and private sector contributions through the Foundation for the National Institutes of Health (FNIH) including generous contributions from the following: AbbVie, Alzheimer’s Association; Alzheimer’s Drug Discovery Foundation; Araclon Biotech; BioClinica, Inc.; Biogen; Bristol-Myers Squibb Company; CereSpir, Inc.; Cogstate; Eisai Inc.; Elan Pharmaceuticals, Inc.; Eli Lilly and Company; EuroImmun; F. Hoffmann-La Roche Ltd and its affiliated company Genentech, Inc.; Fujirebio; GE Healthcare; IXICO Ltd.; Janssen Alzheimer Immunotherapy Research & Development, LLC.; Johnson & Johnson Pharmaceutical Research &Development LLC.; Lumosity; Lundbeck; Merck & Co., Inc.; Meso Scale Diagnostics, LLC.; NeuroRx Research; Neurotrack Technologies; Novartis Pharmaceuticals Corporation; Pfizer Inc.; Piramal Imaging; Servier; Takeda Pharmaceutical Company; and Transition Therapeutics.

## CONSENT STATEMENT

The study protocol was approved by the Nathan S. Kline Institute/Rockland Psychiatric Center Institutional Review Board. All participants provided informed consent.

