## Supplementary Material for "Sex differences in astrocytic reactivity and its association with cognitive decline in cognitively unimpaired older adults"

**Short Title:** Sex Differences in Astrocytic Reactivity and Cognitive Decline

Chelsea Reichert Plaska^a,b^, Tovia Jacobs^c^, Davide Bruno^d,e^, Sang Han Lee^f,g^, Bruno P. Imbimbo^h^, Ricardo Osorio^i,j^, Andréa L. Benedet^k^, Nicholas J. Ashton^l,m,n^, Henrik Zetterberg^o-u^, Nunzio Pomara^v,x^, for the Alzheimer’s Disease Neuroimaging Initiative

**Supplementary Material**

**Supplemental Figure 1. Participant flow diagram**. The total number of ADNI participants considered for this validation analysis is indicated in the top box. Each subsequent box describes the number of ADNI participants, out of the total, with plasma biomarkers, *APOE* genotype and were classified as cognitively unimpaired (CU). The number of ADNI participants included in each of the cross-sectional analyses is described in the bottom boxes for the BBM analysis, global cognition analysis, and the cognitive domain analysis.

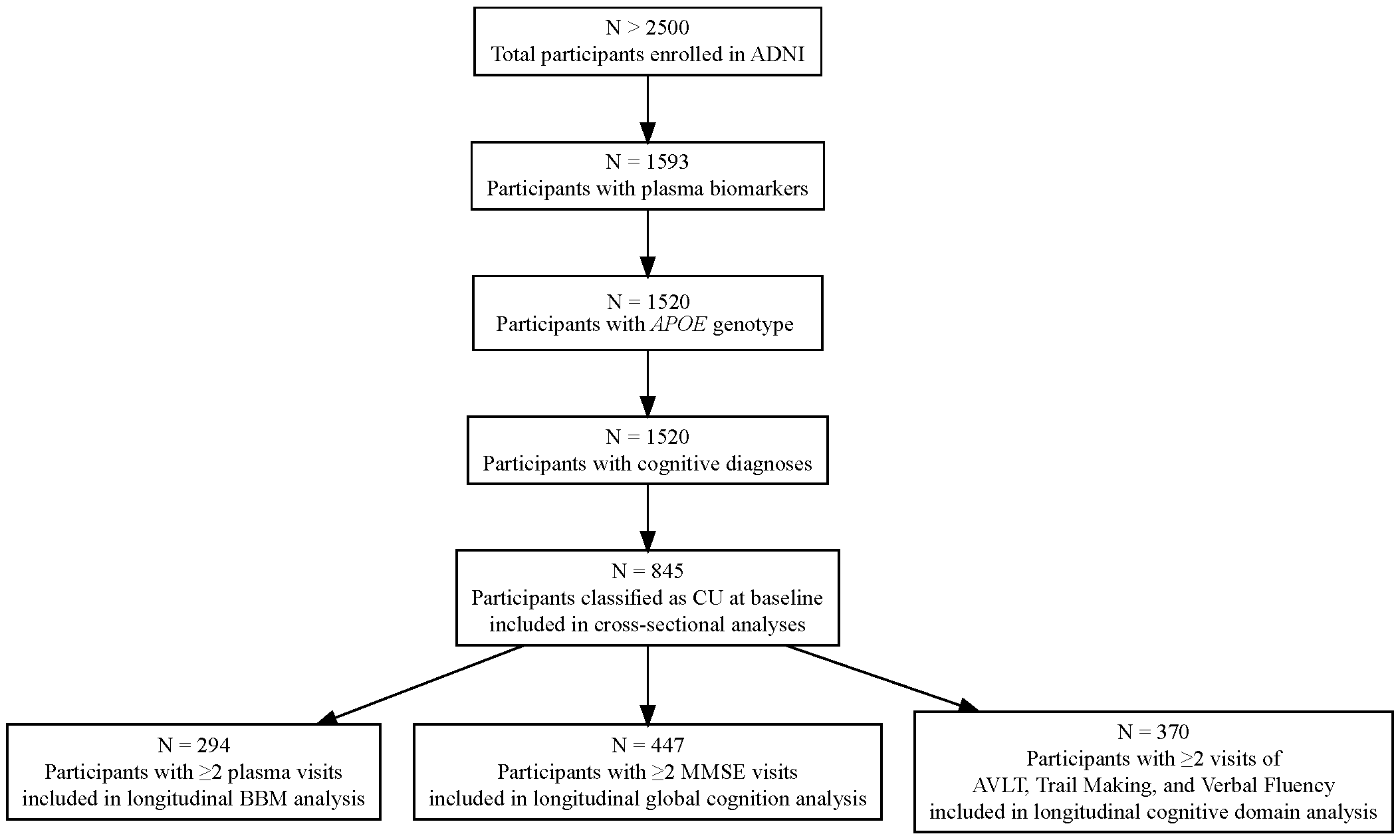

**Supplemental Table 1**. Group comparisons of MERI and ADNI Cohorts by Sex (Female, Male) of performance on select tests from the MERI cognitive battery. The sample is a subset of the whole cohort who were selected as described in Figure 1. Group comparisons by sex were run as independent samples t-test. Significant is p<0.05 and is indicated by an asterisks (*). Mean (SD) reported in the table.

|  | **MERI Cohort** | |  | | **ADNI Cohort** | |  | |
| --- | --- | --- | --- | --- | --- | --- | --- | --- |
|  | **Female** | **Male** | **p** | **Female** | | **Male** | | **p** |
| **n** | 142 | 100 |  | 217 | | 153 | |  |
| **Baseline MMSE**** | 28.99 (1.38) | 28.89 (1.38) | 0.568 | 29.25 (1.04) | | 28.84 (1.27) | | 0.001* |
| **AVLT, total learned** | 47.87 (11.98) | 40.67 (11.73) | <0.001* | 47.79 (10.54) | | 41.09 (9.83) | | <0.001* |
| **AVLT, delayed recall** | 9.04 (4.24) | 7.14 (3.66) | <0.001* | 8.66 (4.04) | | 6.17 (3.76) | | <0.001* |
| **Trail making A, time** | 36.39 (13.06) | 42.14 (21.31) | 0.01* | 30.25 (9.96) | | 36.30 (14.73) | | <0.001* |
| **Tail making B, time** | 96.94 (53.52) | 115.43 (75.05) | 0.026* | 75.91 (36.51) | | 83.64 (38.84) | | 0.052 |
| **Trail making, B-A** | 60.56 (47.33) | 73.29 (60.73) | 0.068 | 45.65 (32.93) | | 47.34 (32.91) | | 0.628 |
| **Trail making, B/A** | 2.70 (1.25) | 2.78 (1.17) | 0.615 | 2.57 (1.07) | | 2.40 (0.84) | | 0.096 |
| **Verbal fluency, total animals** | 19.40 (5.40) | 17.11 (6.24) | 0.003* | 21.60 (5.20) | | 20.41 (5.10) | | 0.028* |
| **Verbal fluency, total F+A+S** | 45.89 (15.00) | 43.44 (14.90) | 0.21 | 15.08 (4.96) | | 14.54 (4.70) | | 0.292 |
| ***MMSE was not completed at the ADNI baseline visit* | | | | | | | | |
| *~Verbal fluency in the ADNI only included total for letter F* | | | | | | | | |

**Supplemental Figure 2. Boxplots of Blood-based Biomarkers (BBMs)**. Boxplots of Baseline ADNI Cohort with individual points superimposed. Comparisons were plotted by sex (female: red, male: blue): Aβ40 (A), Aβ42 (B), Aβ42/40 (C), GFAP (D), p-Tau217 (E), and NfL (F). Significant associations are indicated with an asterisks (p<0.05*, p<0.01**, p<0.001***) and p > 0.05 indicated with ns.

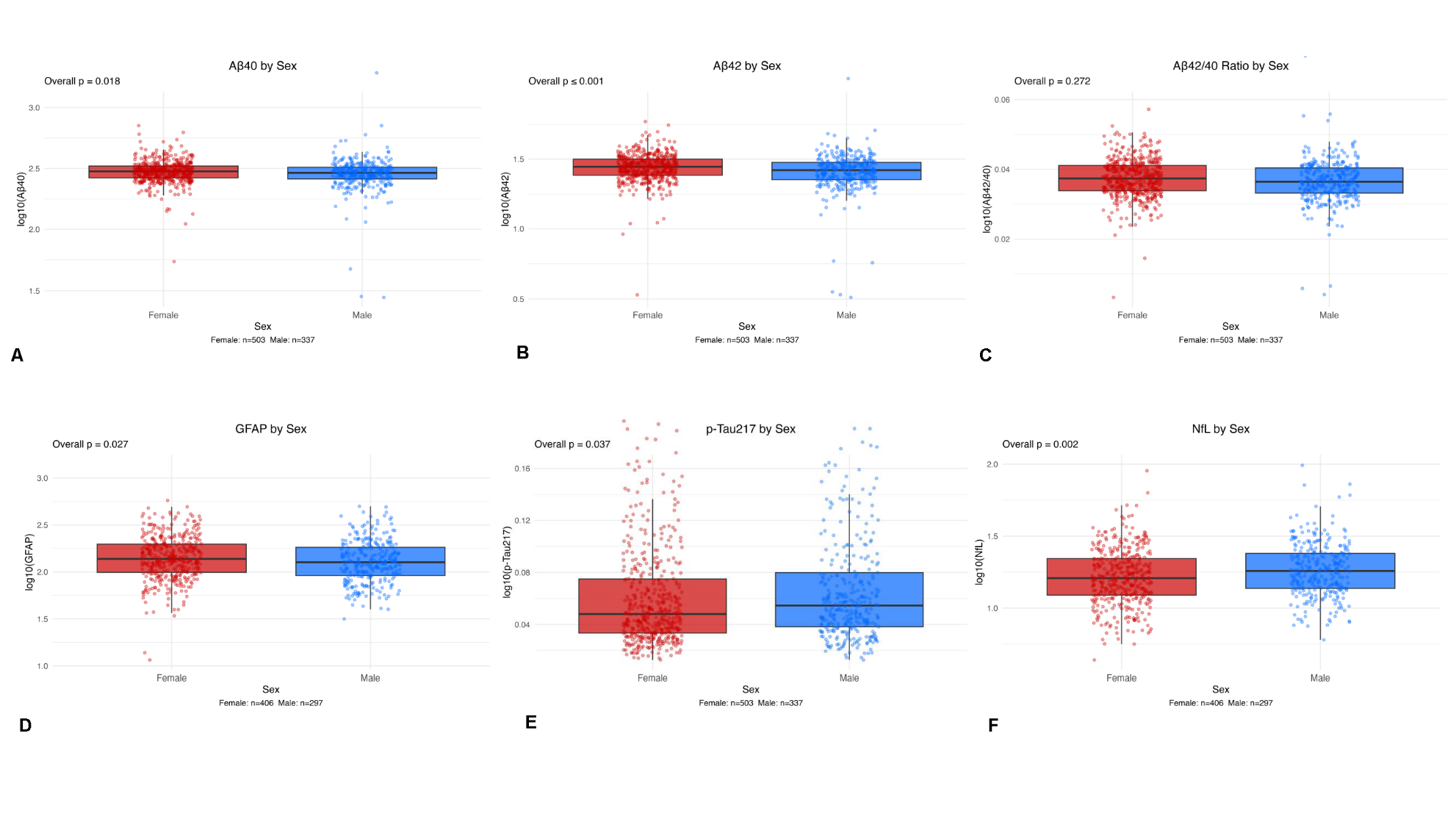

**Supplemental Figure 3.** Heatmaps of Baseline Associations of MERI Cohort for (a) females and (b) males. Red indicates positive association, Blue indicates negative association. Significant associations are indicated with an asterisks (p<0.05*, p<0.01**, p<0.001***).

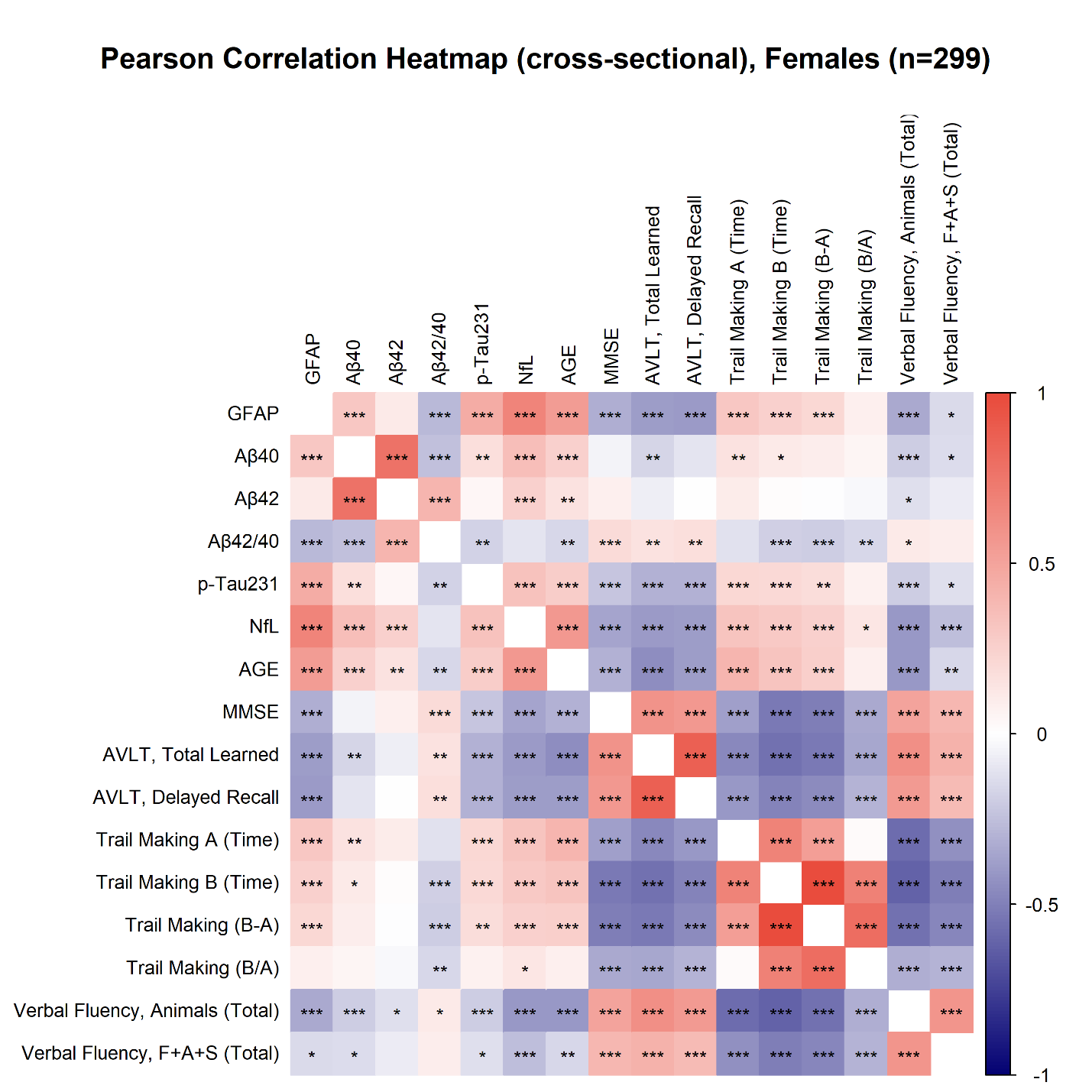

**A**

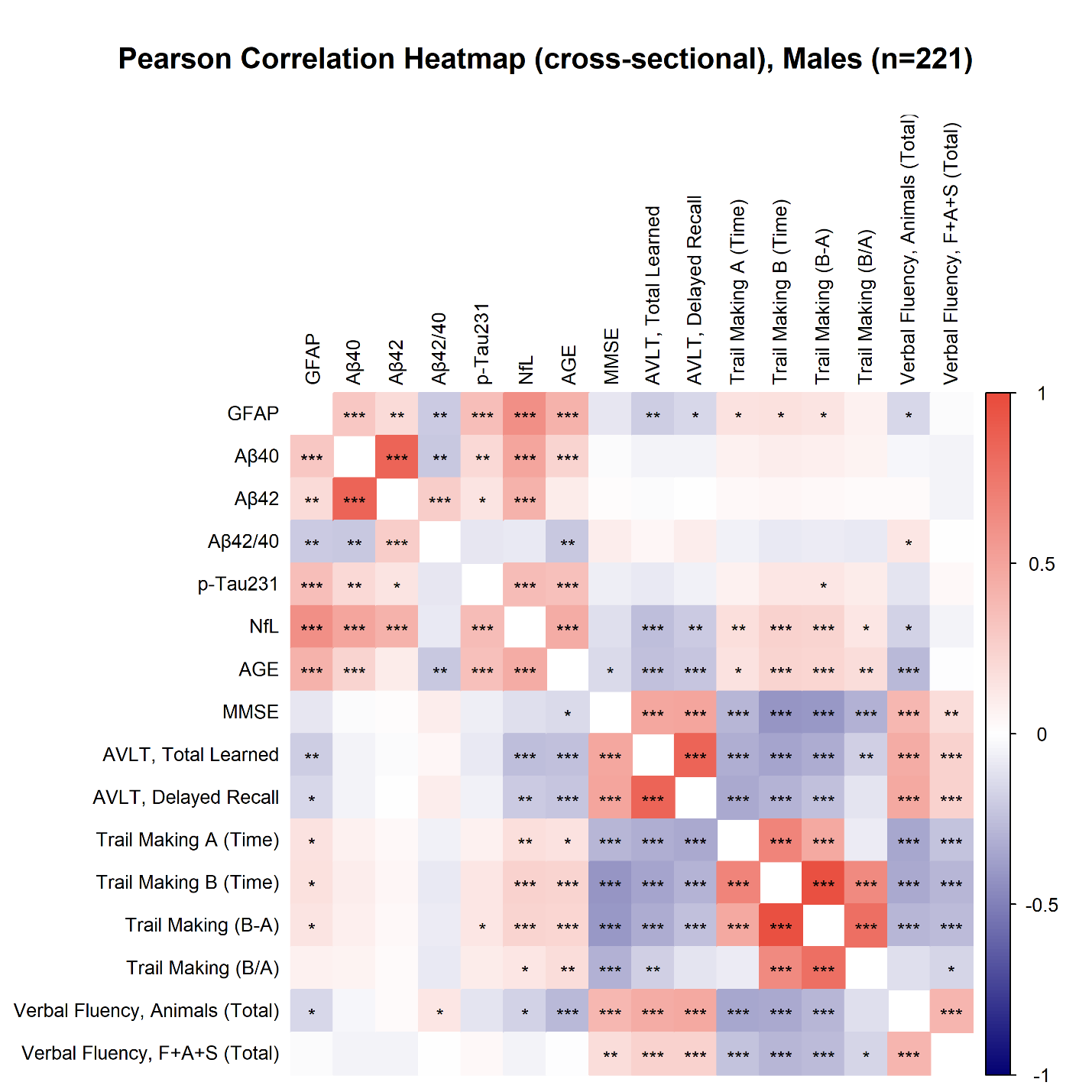

**B**

**Supplemental Figure 4. Longitudinal Aβ42/40 and p-Tau217 trajectories of participants in the lower vs. upper median GFAP at baseline for the ADNI Cohort**. Unadjusted linear trendlines depicting longitudinal plasma obtained throughout follow up visits, comparing participants who were in the lower GFAP median (solid line) versus upper GFAP median (colored shading) at baseline. Individual observations superimposed over trendlines. Comparisons were plotted in the full cohort (A,D), male participants only (B, E), and female participants only (C, F). Plots A-C are for Aβ42/40 ratio and D-F are for p-Tau217. Y-axis is plasma variable and X-axis is time (years). Plots were constructed from the subset of 294 participants with 2+ visits where plasma Aβ42/40/p-Tau217.

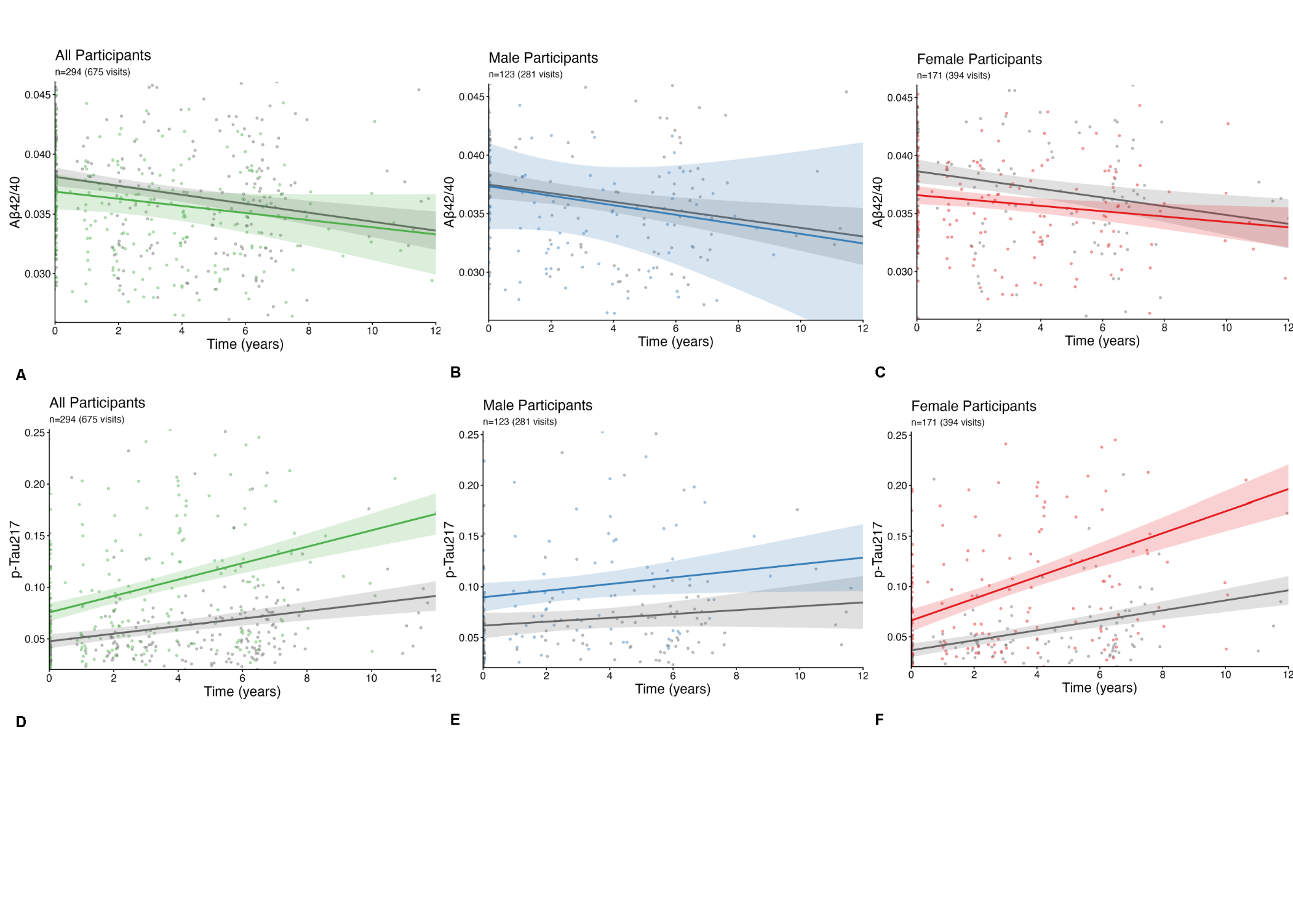

**Supplemental Figure 5.**  **Longitudinal MMSE and AVLT score trajectories of female participants in the lower vs. upper median GFAP at baseline for the ADNI Cohort.** Unadjusted linear trendlines depicting longitudinal MMSE scores obtained throughout follow up visits, comparing participants who were in the lower GFAP median (gray shading) versus upper GFAP median (colored shading) at baseline. Individual observations superimposed over trendlines. Comparisons were plotted in the full cohort (A, D), male participants only (B, E), female participants only (C, F). Plots A-C are for MMSE and D-F are for domain-specific cognition (AVLT Delayed Recall). Plots were constructed from the subset of 294 participants with 2+ visits where MMSE or AVLT were available. Y-axis is total score and X-axis is time (years).

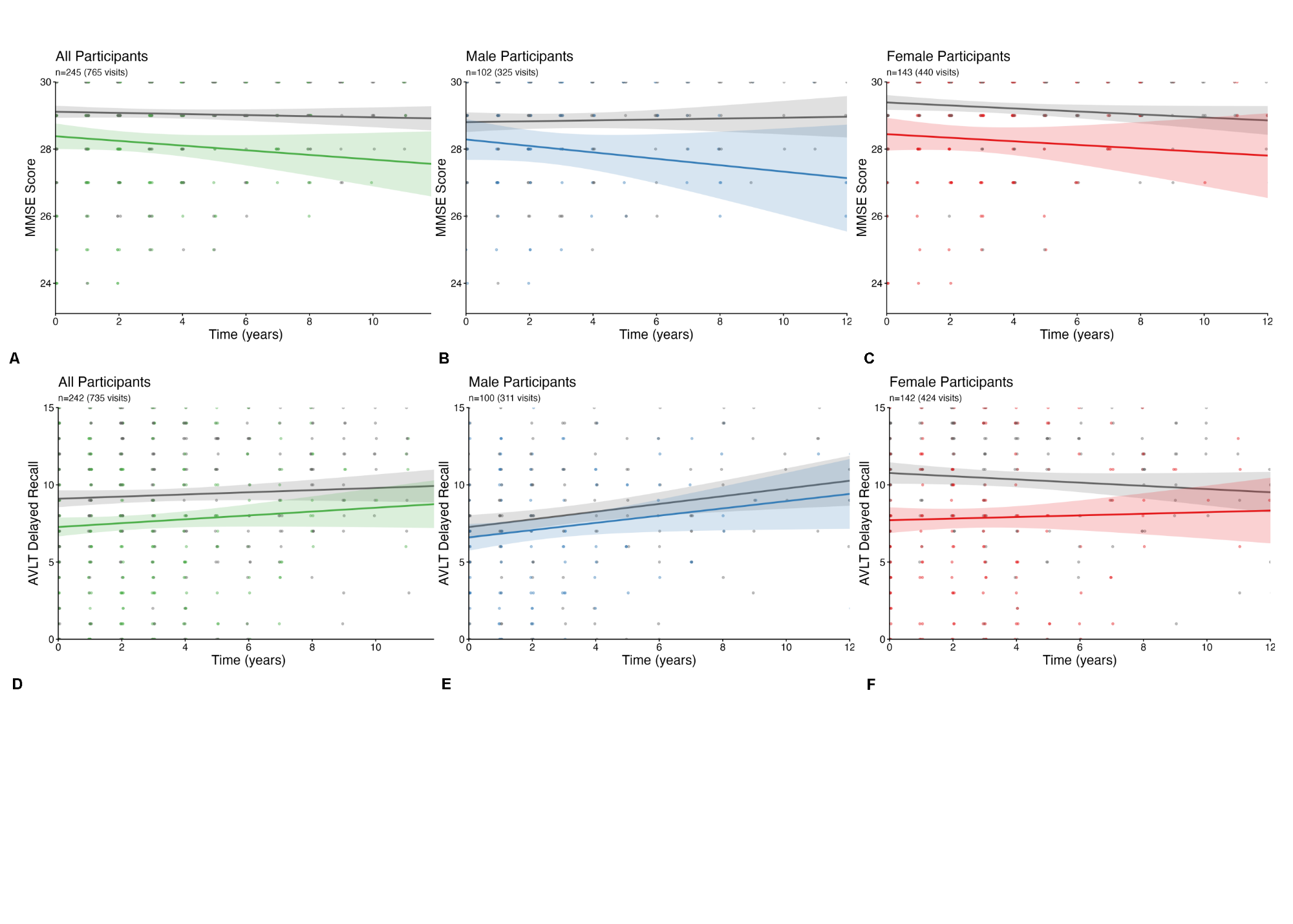

**Supplemental Table 2**. Associations between baseline GFAP and baseline BBMs and cognition in the

the full MERI cohort, in males, and in females. N represents the number of unique participants. Models for the full cohort were adjusted for age at baseline, sex, race, education, and APOE4 status; models for males and females were adjusted for age at baseline, race, education, and APOE4 status. Significant associations are indicated with asterisks (p<0.05*, p<0.01**, p<0.001***). Superscript letters indicate covariates significantly associated with the outcome in that model: a = age at baseline, b = sex, c = race, d = education, e = APOE4; — = no significant covariates.

| **Outcome** | **Full Cohort** | | | **Males** | | | **Females** | | |
| --- | --- | --- | --- | --- | --- | --- | --- | --- | --- |
|  | **N** | **GFAP β (SE)** | **GFAP p-value** | **N** | **GFAP β (SE)** | **GFAP p-value** | **N** | **GFAP β (SE)** | **GFAP p-value** |
| Aβ40 | 581 | 34.656 (7.131) | <0.001*** ^a,b^ | 244 | 38.866 (12.358) | 0.002** ^—^ | 337 | 31.561 (8.218) | <0.001*** ^—^ |
| Aβ42 | 565 | 1.303 (0.423) | 0.002** ^b,e^ | 240 | 1.859 (0.716) | 0.010** ^e^ | 325 | 0.817 (0.502) | 0.105 ^e^ |
| Aβ42/40 | 565 | -0.008 (0.002) | <0.001*** ^a,e^ | 240 | -0.006 (0.003) | 0.111 ^a,e^ | 325 | -0.011 (0.003) | <0.001*** ^d,e^ |
| p-Tau231 | 583 | 0.289 (0.039) | <0.001*** ^a,b,e^ | 247 | 0.222 (0.055) | <0.001*** ^a,e^ | 336 | 0.345 (0.055) | <0.001*** ^e^ |
| NfL | 584 | 0.552 (0.036) | <0.001*** ^a,b^ | 247 | 0.555 (0.056) | <0.001*** ^a,e^ | 337 | 0.539 (0.047) | <0.001*** ^a^ |
| MMSE | 584 | -1.218 (0.305) | <0.001*** ^a,c,d,e^ | 247 | -0.885 (0.447) | 0.049* ^a,c,d,e^ | 337 | -1.476 (0.418) | <0.001*** ^a,c,d^ |
| AVLT, Total Learned | 582 | -9.330 (2.232) | <0.001*** ^a,b,c,d^ | 246 | -7.070 (3.160) | 0.026* ^a,c,d^ | 336 | -10.414 (3.115) | 0.001*** ^a,d^ |
| AVLT, Delayed Recall | 581 | -3.302 (0.763) | <0.001*** ^a,b,c,d^ | 245 | -1.620 (1.047) | 0.123 ^a,c,d^ | 336 | -4.561 (1.088) | <0.001*** ^a,d^ |
| Trail Making A (Time) | 580 | 11.148 (4.487) | 0.013* ^a,b,c,d^ | 247 | 11.894 (7.762) | 0.127 ^—^ | 333 | 8.430 (5.130) | 0.101 ^a,c,d^ |
| Trail Making B (Time) | 544 | 26.066 (12.500) | 0.038* ^a,c,d^ | 231 | 22.420 (18.237) | 0.220 ^a,c,d^ | 313 | 26.600 (17.403) | 0.127 ^a,c,d^ |
| Trail Making (B-A) | 544 | 17.915 (10.822) | 0.098 ^a,c,d^ | 231 | 14.790 (15.279) | 0.334 ^a,c,d^ | 313 | 19.957 (15.443) | 0.197 ^a,c,d^ |
| Trail Making (B/A) | 544 | 0.088 (0.233) | 0.706 ^a,c,d^ | 231 | 0.047 (0.303) | 0.878 ^a,c,d^ | 313 | 0.185 (0.347) | 0.594 ^c,d^ |
| Verbal Fluency, Animals (Total) | 579 | -4.055 (1.075) | <0.001*** ^a,b,c,d^ | 245 | -2.723 (1.657) | 0.102 ^a,c^ | 334 | -5.067 (1.417) | <0.001*** ^a,c,d^ |
| Verbal Fluency, F+A+S (Total) | 581 | -5.697 (2.842) | 0.046* ^b,c,d^ | 246 | -3.972 (4.230) | 0.349 ^c,d^ | 335 | -6.248 (3.841) | 0.105 ^c,d^ |

**Supplemental Table 3**. Associations between baseline GFAP and baseline BBMs and cognition in a) the full MERI cohort and b) the full ADNI cohort. N represents the number of unique participants. Each model was adjusted for age at baseline, sex, race, education, and APOE4 status. Female sex was the reference group; therefore, the GFAP β represents the association among females, and the GFAP × sex β represents the difference in this association for males relative to females. Significant associations are indicated with asterisks (p<0.05*, p<0.01**, p<0.001***).

| **Outcome** | **N** | **GFAP β (SE)** | **GFAP p-value** | **GFAP*Sex Beta (SE)** | **GFAP*Sex p-value** | **Significant Covariates** |
| --- | --- | --- | --- | --- | --- | --- |
| **a) MERI Cohort** | | | | | | |
| Aβ40 | 581 | 29.382 (9.089) | 0.001*** | 11.625 (12.417) | 0.350 | age at baseline |
| Aβ42 | 565 | 0.970 (0.539) | 0.073 | 0.736 (0.740) | 0.320 | APOE4 |
| Aβ42/40 | 565 | -0.009 (0.003) | 0.002** | 0.002 (0.004) | 0.629 | age at baseline, APOE4 |
| p-Tau231 | 583 | 0.326 (0.050) | <0.001*** | -0.081 (0.068) | 0.230 | age at baseline, APOE4 |
| NfL | 584 | 0.558 (0.046) | <0.001*** | -0.014 (0.062) | 0.818 | age at baseline |
| MMSE | 584 | -1.590 (0.389) | <0.001*** | 0.816 (0.530) | 0.124 | age at baseline, race other, education, APOE4 |
| AVLT, Total Learned | 582 | -12.980 (2.844) | <0.001*** | 7.985 (3.873) | 0.040* | age at baseline, race, education, sex |
| AVLT, Delayed Recall | 581 | -5.037 (0.967) | <0.001*** | 3.813 (1.319) | 0.004** | age at baseline, race, education, sex |
| Trail Making A (Time) | 580 | 14.249 (5.734) | 0.013* | -6.778 (7.799) | 0.385 | age at baseline, race, education |
| Trail Making B (Time) | 544 | 32.459 (15.959) | 0.042* | -14.046 (21.778) | 0.519 | age at baseline, race, education |
| Trail Making (B-A) | 544 | 20.946 (13.820) | 0.130 | -6.657 (18.860) | 0.724 | age at baseline, race, education |
| Trail Making (B/A) | 544 | 0.022 (0.298) | 0.941 | 0.145 (0.406) | 0.722 | age at baseline, race, education |
| Verbal Fluency, Animals (Total) | 579 | -5.327 (1.373) | <0.001*** | 2.777 (1.868) | 0.138 | age at baseline, race, education |
| Verbal Fluency, F+A+S (Total) | 581 | -8.504 (3.628) | 0.019* | 6.157 (4.948) | 0.214 | race, education |
| **b) ADNI Cohort** | | | | | | |
| Aβ40 | 707 | 0.032 (0.026) | 0.220 | 0.059 (0.038) | 0.122 | age at baseline |
| Aβ42 | 707 | -0.006 (0.026) | 0.820 | 0.045 (0.038) | 0.237 | age at baseline, APOE4 |
| Aβ42/40 | 707 | -0.003 (0.002) | 0.188 | -0.000 (0.003) | 0.945 | APOE4 |
| p-Tau217 | 707 | 0.068 (0.011) | <0.001*** | 0.035 (0.016) | 0.028* | age at baseline, race, APOE4 |
| NfL | 707 | 0.337 (0.033) | <0.001*** | 0.013 (0.047) | 0.790 | age at baseline, race, APOE4 |
| MMSE | 622 | -0.265 (0.303) | 0.383 | 0.070 (0.450) | 0.877 | age at baseline, race, education |
| AVLT, Total Learned | 622 | 0.921 (2.454) | 0.708 | -0.948 (3.637) | 0.794 | age at baseline, education |
| AVLT, Delayed Recall | 622 | 0.149 (0.974) | 0.878 | -0.995 (1.444) | 0.491 | age at baseline, race, education |
| Trail Making A (Time) | 622 | 3.071 (2.791) | 0.272 | 1.167 (4.137) | 0.778 | age at baseline, race |
| Trail Making B (Time) | 622 | 16.298 (8.966) | 0.070 | -21.683 (13.292) | 0.103 | age at baseline, race, education |
| Verbal Fluency, Animals (Total) | 622 | -0.620 (1.225) | 0.613 | -0.722 (1.817) | 0.691 | age at baseline, race, education |
| Verbal Fluency, F (Total) | 622 | -0.726 (1.143) | 0.526 | 0.896 (1.694) | 0.597 | education |

**Supplemental Table 4**. Associations between baseline GFAP and baseline BBMs and cognition in the full ADNI cohort, in males, and in females. N represents the number of unique participants. Models for the full cohort were adjusted for age at baseline, sex, race, education, and APOE4 status; models for males and females were adjusted for age at baseline, race, education, and APOE4 status. Significant associations are indicated with asterisks (p<0.05*, p<0.01**, p<0.001***). Superscript letters indicate covariates significantly associated with the outcome in that model: a = age at baseline, b = sex, c = race, d = education, e = APOE4; — = no significant covariates.

| **Outcome** | **Full Cohort** | | | **Males** | | | **Females** | | |
| --- | --- | --- | --- | --- | --- | --- | --- | --- | --- |
|  | **N** | **GFAP β (SE)** | **GFAP p-value** | **N** | **GFAP β (SE)** | **GFAP p-value** | **N** | **GFAP β (SE)** | **GFAP p-value** |
| Aβ40 | 707 | 0.056 (0.021) | 0.008** ^a,b^ | 299 | 0.065 (0.040) | 0.104 ^a^ | 408 | 0.049 (0.023) | 0.029* ^a,e^ |
| Aβ42 | 707 | 0.012 (0.021) | 0.559 ^a,b,e^ | 299 | 0.003 (0.041) | 0.935 ^a,e^ | 408 | 0.018 (0.021) | 0.383 ^e^ |
| Aβ42/40 | 707 | -0.003 (0.002) | 0.093 ^e^ | 299 | -0.003 (0.004) | 0.335 ^e^ | 408 | -0.002 (0.002) | 0.124 ^a,e^ |
| p-Tau217 | 707 | 0.082 (0.009) | <0.001*** ^a,b,c,e^ | 299 | 0.105 (0.015) | <0.001*** ^c^ | 408 | 0.068 (0.011) | <0.001*** ^a,e^ |
| NfL | 707 | 0.342 (0.026) | <0.001*** ^a,c,e^ | 299 | 0.347 (0.041) | <0.001*** ^a,c,d^ | 408 | 0.335 (0.034) | <0.001*** ^a,c^ |
| MMSE | 622 | -0.238 (0.250) | 0.341 ^a,b,c,d^ | 252 | 0.118 (0.482) | 0.807 ^a,d^ | 370 | -0.482 (0.270) | 0.075 ^c,d^ |
| AVLT, Total Learned | 622 | 0.559 (2.021) | 0.782 ^a,b,d^ | 252 | -0.487 (3.333) | 0.884 ^a,d^ | 370 | 0.736 (2.556) | 0.774 ^a,c,d^ |
| AVLT, Delayed Recall | 622 | -0.231 (0.803) | 0.774 ^a,b,c,d^ | 252 | -0.958 (1.301) | 0.462 ^d^ | 370 | -0.064 (1.024) | 0.950 ^a,c,d^ |
| Trail Making A (Time) | 622 | 3.517 (2.299) | 0.127 ^a,b,c^ | 252 | 0.675 (4.301) | 0.875 ^a,c^ | 370 | 4.392 (2.584) | 0.090 ^a,c^ |
| Trail Making B (Time) | 622 | 8.020 (7.402) | 0.279 ^a,c,d^ | 252 | -6.724 (11.670) | 0.565 ^a,c,d^ | 370 | 16.873 (9.681) | 0.082 ^a,c,d^ |
| Verbal Fluency, Animals (Total) | 622 | -0.895 (1.010) | 0.376 ^a,b,c,d^ | 252 | -1.089 (1.599) | 0.497 ^a,c,d^ | 370 | -0.610 (1.311) | 0.642 ^a,c,d^ |
| Verbal Fluency, F (Total) | 622 | -0.384 (0.942) | 0.684 ^d^ | 252 | -0.691 (1.527) | 0.651 ^d^ | 370 | -0.269 (1.211) | 0.825 ^d^ |

**Supplemental Table 5.** Associations between baseline GFAP and longitudinal BBMs in the full MERI cohort, including three- way interactions between baseline GFAP, sex, and time since baseline. Output of linear mixed effects models. Females are the reference group for sex interactions. Each model was adjusted for age at baseline, race, education, and APOE4 status, with time since baseline included as part of the interaction model. Significant associations are indicated with asterisks (p<0.05*, p<0.01**, p<0.001***).

| **Outcome (longitudinal)** | **GFAP** | | **GFAP × Time** | | **GFAP × Sex** | | **Sex × Time** | | **GFAP × Sex × Time** | | **Significant Covariates** |
| --- | --- | --- | --- | --- | --- | --- | --- | --- | --- | --- | --- |
|  | **β (SE)** | **p-value** | **β (SE)** | **p-value** | **β (SE)** | **p-value** | **β (SE)** | **p-value** | **β (SE)** | **p-value** |  |
| Aβ40 | 17.181 (11.746) | 0.145 | 1.670 (2.138) | 0.435 | 11.970 (15.839) | 0.450 | 9.127 (7.091) | 0.199 | -5.081 (3.424) | 0.138 | age at baseline |
| Aβ42 | -0.213 (0.762) | 0.780 | 0.090 (0.124) | 0.470 | 1.235 (1.026) | 0.230 | 0.384 (0.412) | 0.351 | -0.213 (0.199) | 0.284 | age at baseline, education, APOE4 |
| Aβ42/40 | -0.012 (0.005) | 0.012* | 0.000 (0.001) | 0.782 | 0.005 (0.006) | 0.392 | -0.002 (0.002) | 0.151 | 0.001 (0.001) | 0.121 | APOE4 |
| p-Tau231 | 0.236 (0.070) | 0.001*** | -0.010 (0.011) | 0.335 | -0.128 (0.094) | 0.176 | -0.036 (0.035) | 0.303 | 0.020 (0.017) | 0.236 | age at baseline, APOE4 |
| NfL | 0.368 (0.080) | <0.001*** | -0.008 (0.011) | 0.448 | 0.035 (0.107) | 0.743 | 0.028 (0.035) | 0.420 | -0.013 (0.017) | 0.443 | age at baseline, race |

**Supplemental Table 5b**. Associations between baseline GFAP and longitudinal blood-based biomarkers for the interaction between baseline GFAP and time, in the full MERI cohort, males only, and females only. Output of linear mixed effects models. Full-cohort models were adjusted for age at baseline, time since baseline, race, education, sex, and APOE4 status; male- and female-only models were adjusted for age at baseline, time since baseline, race, education, and APOE4 status. Significant associations are indicated with asterisks (p<0.05*, p<0.01**, p<0.001***). Superscript letters on the GFAP × Time p-value indicate covariates significantly associated with the outcome in that model: a = age at baseline, b = sex, c = race, d = education, e = APOE4, f = time since baseline, g = time since baseline:GFAP interaction; — = no significant covariates.

| **Outcome (longitudinal)** | **Full Cohort** | | | | **Males** | | | | **Females** | | | |
| --- | --- | --- | --- | --- | --- | --- | --- | --- | --- | --- | --- | --- |
|  | **GFAP β (SE)** | **GFAP p-value** | **GFAP × Time β (SE)** | **GFAP × Time p-value** | **GFAP β (SE)** | **GFAP p-value** | **GFAP × Time β (SE)** | **GFAP × Time p-value** | **GFAP β (SE)** | **GFAP p-value** | **GFAP × Time β (SE)** | **GFAP × Time p-value** |
| Aβ40 | 21.668 (8.526) | 0.012 ^—^ | 0.294 (1.631) | 0.857 ^a^ | 20.664 (13.767) | 0.136 ^—^ | -3.562 (2.856) | 0.214 ^a^ | 26.053 (10.767) | 0.017* ^—^ | 1.743 (2.002) | 0.385 ^—^ |
| Aβ42 | 0.357 (0.555) | 0.521 ^—^ | 0.029 (0.095) | 0.763 ^a,d,e^ | 0.981 (0.873) | 0.264 ^—^ | -0.123 (0.158) | 0.439 ^—^ | -0.291 (0.740) | 0.695 ^—^ | 0.086 (0.122) | 0.482 ^a,e^ |
| Aβ42/40 | -0.009 (0.003) | 0.011* ^—^ | 0.001 (0.000) | 0.160 ^e^ | -0.002 (0.005) | 0.599 ^—^ | 0.001 (0.001) | 0.042* ^a,e,f,g^ | -0.017 (0.005) | 0.001*** ^—^ | 0.000 (0.000) | 0.762 ^e^ |
| p-Tau231 | 0.177 (0.051) | 0.001*** ^—^ | -0.004 (0.008) | 0.585 ^a,b,e^ | 0.087 (0.074) | 0.245 ^—^ | 0.010 (0.012) | 0.403 ^a^ | 0.259 (0.073) | 0.001*** ^—^ | -0.010 (0.011) | 0.367 ^e^ |
| NfL | 0.385 (0.058) | <0.001*** ^—^ | -0.014 (0.008) | 0.087 ^a,c,f^ | 0.397 (0.092) | <0.001*** ^—^ | -0.021 (0.015) | 0.176 ^a,f^ | 0.353 (0.079) | <0.001*** ^—^ | -0.008 (0.009) | 0.374 ^a^ |

**Supplemental Table 6.** Associations between baseline GFAP and longitudinal BBMs in a) the whole ADNI cohort, b) males only, and c) females only. Output of linear mixed effects models. Each model was adjusted for age at baseline, time since baseline, race, education, and APOE4 status (sex was additionally adjusted for in the whole-cohort model). Significant associations are indicated with asterisks (p<0.05*, p<0.01**, p<0.001***).

| **Outcome (longitudinal)** | **Baseline GFAP, β (SE)** | **Baseline GFAP, p-value** | **Significant Covariates** |
| --- | --- | --- | --- |
| **a) Whole ADNI Cohort (N=294 participants across 675 total visits)** | | | |
| Aβ40 | 0.047 (0.023) | 0.045* | age at baseline, education, APOE4, time since baseline |
| Aβ42 | 0.010 (0.021) | 0.648 | age at baseline, education, APOE4, time since baseline |
| Aβ42/40 | -0.003 (0.002) | 0.119 | APOE4, time since baseline |
| p-Tau217 | 0.092 (0.014) | <0.001*** | age at baseline, race, APOE4, time since baseline |
| NfL | 0.320 (0.041) | <0.001*** | age at baseline, race, time since baseline |
| **b) Males Only (N=123 participants across 281 total visits)** | | | |
| Aβ40 | 0.003 (0.041) | 0.944 | age at baseline, time since baseline |
| Aβ42 | -0.026 (0.033) | 0.435 | age at baseline, APOE4, time since baseline |
| Aβ42/40 | -0.002 (0.004) | 0.560 | APOE4, time since baseline |
| p-Tau217 | 0.105 (0.024) | <0.001*** | time since baseline |
| NfL | 0.380 (0.069) | <0.001*** | age at baseline, time since baseline |
| **c) Females Only (N=171 participants across 394 total visits)** | | | |
| Aβ40 | 0.077 (0.027) | 0.005** | education, APOE4, time since baseline |
| Aβ42 | 0.036 (0.027) | 0.175 | education, APOE4, time since baseline |
| Aβ42/40 | -0.003 (0.001) | 0.025* | age at baseline, APOE4, time since baseline |
| p-Tau217 | 0.082 (0.017) | <0.001*** | age at baseline, race, APOE4, time since baseline |
| NfL | 0.289 (0.050) | <0.001*** | age at baseline, education, APOE4, time since baseline |

**Supplemental Table 7.** Associations between baseline GFAP and longitudinal BBMs in the full ADNI cohort, including three-way interactions between baseline GFAP, sex, and time since baseline. Output of linear mixed effects models. Females are the reference group for sex interactions. Each model was adjusted for age at baseline, race, education, and APOE4 status, with time since baseline included as part of the interaction model. Significant associations are indicated with asterisks (p<0.05*, p<0.01**, p<0.001***).

| **Outcome (longitudinal)** | **GFAP** | | **GFAP × Time** | | **GFAP × Sex** | | **Sex × Time** | | **GFAP × Sex × Time** | | **Significant Covariates** |
| --- | --- | --- | --- | --- | --- | --- | --- | --- | --- | --- | --- |
|  | **β (SE)** | **p-value** | **β (SE)** | **p-value** | **β (SE)** | **p-value** | **β (SE)** | **p-value** | **β (SE)** | **p-value** |  |
| Aβ40 | 0.061 (0.034) | 0.070 | -0.001 (0.007) | 0.840 | -0.036 (0.050) | 0.466 | -0.002 (0.019) | 0.926 | 0.004 (0.009) | 0.685 | age at baseline, education, APOE4 |
| Aβ42 | 0.017 (0.029) | 0.573 | 0.002 (0.005) | 0.772 | -0.013 (0.044) | 0.765 | 0.013 (0.015) | 0.383 | -0.004 (0.007) | 0.579 | age at baseline, education, APOE4 |
| Aβ42/40 | -0.004 (0.003) | 0.160 | 0.000 (0.001) | 0.556 | 0.003 (0.004) | 0.469 | 0.002 (0.002) | 0.330 | -0.001 (0.001) | 0.276 | APOE4 |
| p-Tau217 | 0.067 (0.019) | <0.001*** | 0.007 (0.003) | 0.017* | 0.032 (0.028) | 0.245 | 0.009 (0.008) | 0.238 | -0.006 (0.004) | 0.102 | age at baseline, race, APOE4 |
| NfL | 0.294 (0.055) | <0.001*** | -0.007 (0.010) | 0.497 | 0.098 (0.081) | 0.229 | -0.007 (0.028) | 0.803 | 0.003 (0.014) | 0.827 | age at baseline, race |

**Supplemental Table 7b**. Associations between baseline GFAP and longitudinal blood-based biomarkers for the interaction between baseline GFAP and time, in the full ADNI cohort, males only, and females only. Output of linear mixed effects models. Full-cohort models were adjusted for age at baseline, time since baseline, race, education, sex, and APOE4 status; male- and female-only models were adjusted for age at baseline, time since baseline, race, education, and APOE4 status. Significant associations are indicated with asterisks (p<0.05*, p<0.01**, p<0.001***). Superscript letters on the GFAP × Time p-value indicate covariates significantly associated with the outcome in that model: a = age at baseline, b = sex, c = race, d = education, e = APOE4, f = time since baseline, g = time since baseline:GFAP interaction; — = no significant covariates.

| **Outcome (longitudinal)** | **Full Cohort** | | | | **Males** | | | | **Females** | | | |
| --- | --- | --- | --- | --- | --- | --- | --- | --- | --- | --- | --- | --- |
|  | **GFAP β (SE)** | **GFAP p-value** | **GFAP × Time β (SE)** | **GFAP × Time p-value** | **GFAP β (SE)** | **GFAP p-value** | **GFAP × Time β (SE)** | **GFAP × Time p-value** | **GFAP β (SE)** | **GFAP p-value** | **GFAP × Time β (SE)** | **GFAP × Time p-value** |
| Aβ40 | 0.052 (0.027) | 0.053 ^—^ | -0.002 (0.004) | 0.701 ^a,d,e^ | -0.005 (0.048) | 0.917 ^—^ | 0.002 (0.007) | 0.750 ^a^ | 0.084 (0.030) | 0.006** ^—^ | -0.003 (0.005) | 0.632 ^d,e^ |
| Aβ42 | 0.017 (0.023) | 0.471 ^—^ | -0.002 (0.004) | 0.490 ^a,d,e^ | -0.020 (0.038) | 0.602 ^—^ | -0.002 (0.005) | 0.733 ^a,e^ | 0.034 (0.030) | 0.250 ^—^ | 0.001 (0.005) | 0.874 ^d,e^ |
| Aβ42/40 | -0.003 (0.002) | 0.213 ^—^ | -0.000 (0.000) | 0.917 ^e^ | -0.001 (0.005) | 0.889 ^—^ | -0.000 (0.001) | 0.536 ^e^ | -0.004 (0.002) | 0.013* ^—^ | 0.000 (0.000) | 0.278 ^a,e^ |
| p-Tau217 | 0.078 (0.015) | <0.001*** ^—^ | 0.005 (0.002) | 0.015* ^a,c,e,g^ | 0.104 (0.026) | <0.001*** ^—^ | 0.000 (0.003) | 0.924 ^—^ | 0.064 (0.018) | 0.001*** ^—^ | 0.007 (0.002) | 0.008** ^a,c,e,g^ |
| NfL | 0.330 (0.044) | <0.001*** ^—^ | -0.004 (0.007) | 0.506 ^a,c^ | 0.388 (0.073) | <0.001*** ^—^ | -0.003 (0.009) | 0.756 ^a^ | 0.304 (0.055) | <0.001*** ^—^ | -0.007 (0.010) | 0.513 ^a,d,e^ |

**Supplemental Table 8.** Associations between baseline GFAP and longitudinal MMSE in the MERI cohort, including three-way interactions between baseline GFAP, sex, and time since baseline. Output of linear mixed effects models. Females are the reference group for sex interactions. Each model was adjusted for age at baseline, race, education, and APOE4 status, with time since baseline included as part of the interaction model. Significant associations are indicated with asterisks (p<0.05*, p<0.01**, p<0.001***).

| **Outcome (longitudinal)** | **GFAP** | | **GFAP × Time** | | **GFAP × Sex** | | **Sex × Time** | | **GFAP × Sex × Time** | | **Significant Covariates** |
| --- | --- | --- | --- | --- | --- | --- | --- | --- | --- | --- | --- |
|  | **β (SE)** | **p-value** | **β (SE)** | **p-value** | **β (SE)** | **p-value** | **β (SE)** | **p-value** | **β (SE)** | **p-value** |  |
| MMSE | -1.543 (0.836) | 0.066 | -0.323 (0.109) | 0.003** | 1.010 (1.115) | 0.366 | 0.338 (0.379) | 0.374 | -0.207 (0.185) | 0.263 | race, education |

**Supplemental Table 8b**. Associations between baseline GFAP and longitudinal MMSE for the interaction between baseline GFAP and time, in the full MERI cohort, males only, and females only. Output of linear mixed effects models. Full-cohort models were adjusted for age at baseline, sex, race, education, and APOE4 status; male- and female-only models were adjusted for age at baseline, race, education, and APOE4 status. Significant associations are indicated with asterisks (p<0.05*, p<0.01**, p<0.001***). Superscript letters on the GFAP × Time p-value indicate covariates significantly associated with the outcome in that model: a = age at baseline, b = sex, c = race, d = education, e = APOE4, f = time since baseline, g = time since baseline:GFAP interaction; — = no significant covariates.

| **Outcome (longitudinal)** | **Full Cohort** | | | | **Males** | | | | **Females** | | | |
| --- | --- | --- | --- | --- | --- | --- | --- | --- | --- | --- | --- | --- |
|  | **GFAP β (SE)** | **GFAP p-value** | **GFAP × Time β (SE)** | **GFAP × Time p-value** | **GFAP β (SE)** | **GFAP p-value** | **GFAP × Time β (SE)** | **GFAP × Time p-value** | **GFAP β (SE)** | **GFAP p-value** | **GFAP × Time β (SE)** | **GFAP × Time p-value** |
| MMSE | -1.129 (0.609) | 0.065 ^—^ | -0.359 (0.086) | <0.001*** ^a,b,c,d,f,g^ | -0.463 (0.883) | 0.601 ^—^ | -0.534 (0.151) | <0.001*** ^f,g^ | -1.743 (0.886) | 0.051 ^—^ | -0.321 (0.108) | <0.003** ^f,g^ |

**Supplemental Table 8c**. Associations between baseline GFAP and longitudinal cognition in the MERI cohort, including three-way interactions between baseline GFAP, sex, and time since baseline. Output of linear mixed effects models. Females are the reference group for sex interactions. Each model was adjusted for age at baseline, race, education, and APOE4 status, with time since baseline included as part of the interaction model. Significant associations are indicated with asterisks (p<0.05*, p<0.01**, p<0.001***).

| **Outcome (longitudinal)** | **GFAP** | | **GFAP × Time** | | **GFAP × Sex** | | **Sex × Time** | | **GFAP × Sex × Time** | | **Significant Covariates** |
| --- | --- | --- | --- | --- | --- | --- | --- | --- | --- | --- | --- |
|  | **β (SE)** | **p-value** | **β (SE)** | **p-value** | **β (SE)** | **p-value** | **β (SE)** | **p-value** | **β (SE)** | **p-value** |  |
| AVLT, Total Learned | -10.678 (4.598) | 0.021* | -1.144 (0.447) | 0.011* | 6.322 (6.191) | 0.308 | -1.875 (1.579) | 0.236 | 0.865 (0.773) | 0.263 | age at baseline, race, education |
| AVLT, Delayed Recall | -5.113 (1.639) | 0.002** | -0.512 (0.149) | 0.001*** | 4.655 (2.206) | 0.036* | -0.084 (0.527) | 0.874 | 0.050 (0.258) | 0.845 | age at baseline |
| Trail Making A (Time) | 1.425 (6.144) | 0.817 | 0.792 (0.649) | 0.223 | 1.442 (8.273) | 0.862 | -0.576 (2.289) | 0.801 | 0.333 (1.120) | 0.766 | age at baseline, race |
| Trail Making B (Time) | -12.987 (23.624) | 0.583 | 5.238 (2.716) | 0.054 | 40.582 (31.821) | 0.203 | -6.140 (9.589) | 0.522 | 3.416 (4.692) | 0.467 | age at baseline, race, education |
| Trail Making (B-A) | -13.299 (19.683) | 0.500 | 3.915 (2.649) | 0.140 | 37.525 (26.537) | 0.159 | -5.748 (9.349) | 0.539 | 3.079 (4.574) | 0.501 | age at baseline, race, education |
| Trail Making (B/A) | -0.235 (0.433) | 0.589 | 0.010 (0.071) | 0.888 | 0.453 (0.586) | 0.439 | -0.254 (0.249) | 0.308 | 0.124 (0.122) | 0.310 | age at baseline, race, education |
| Verbal Fluency, Animals (Total) | -3.499 (2.117) | 0.100 | -0.076 (0.262) | 0.771 | 3.525 (2.853) | 0.218 | 0.491 (0.925) | 0.596 | -0.176 (0.453) | 0.697 | age at baseline, race |
| Verbal Fluency, F+A+S (Total) | -7.892 (5.441) | 0.148 | 0.261 (0.501) | 0.602 | 10.958 (7.325) | 0.136 | 0.837 (1.768) | 0.636 | -0.362 (0.865) | 0.676 | race, education |

**Supplemental Table 8d**. Associations between baseline GFAP and longitudinal cognition for the interaction between baseline GFAP and time, in the full MERI cohort, males only, and females only. Output of linear mixed effects models. Full-cohort models were adjusted for age at baseline, sex, race, education, and APOE4 status; male- and female-only models were adjusted for age at baseline, race, education, and APOE4 status. Significant associations are indicated with asterisks (p<0.05*, p<0.01**, p<0.001***). Superscript letters on the GFAP × Time p-value indicate covariates significantly associated with the outcome in that model: a = age at baseline, b = sex, c = race, d = education, e = APOE4, f = time since baseline, g = time since baseline:GFAP interaction; — = no significant covariates.

| **Outcome (longitudinal)** | **Full Cohort** | | | | **Males** | | | | **Females** | | | |
| --- | --- | --- | --- | --- | --- | --- | --- | --- | --- | --- | --- | --- |
|  | **GFAP β (SE)** | **GFAP p-value** | **GFAP × Time β (SE)** | **GFAP × Time p-value** | **GFAP β (SE)** | **GFAP p-value** | **GFAP × Time β (SE)** | **GFAP × Time p-value** | **GFAP β (SE)** | **GFAP p-value** | **GFAP × Time β (SE)** | **GFAP × Time p-value** |
| AVLT, Total Learned | -7.477 (3.395) | 0.029* ^—^ | -0.821 (0.356) | 0.021* ^a,b,c,d,g^ | -4.660 (4.921) | 0.346 ^—^ | -0.263 (0.664) | 0.692 ^—^ | -9.970 (4.872) | 0.042* ^—^ | -1.172 (0.429) | 0.007** ^a,f,g^ |
| AVLT, Delayed Recall | -2.753 (1.221) | 0.025* ^—^ | -0.515 (0.118) | <0.001*** ^a,b,f,g^ | -0.469 (1.652) | 0.777 ^—^ | -0.459 (0.217) | 0.035* ^f,g^ | -5.191 (1.814) | 0.005** ^—^ | -0.526 (0.146) | <0.001*** ^f,g^ |
| Trail Making A (Time) | 2.282 (4.518) | 0.614 ^—^ | 0.852 (0.514) | 0.098 ^a,b,c^ | 5.965 (8.336) | 0.476 ^—^ | 1.215 (0.939) | 0.197 ^—^ | -0.715 (5.124) | 0.889 ^—^ | 0.954 (0.614) | 0.121 ^a,c,e^ |
| Trail Making B (Time) | 8.720 (17.403) | 0.617 ^—^ | 5.841 (2.157) | 0.007** ^a,b,c,d,g^ | 26.900 (31.842) | 0.400 ^—^ | 9.165 (4.280) | 0.033* ^a,g^ | -6.601 (19.080) | 0.730 ^—^ | 4.834 (2.413) | 0.046* ^a,c,d,g^ |
| Trail Making (B-A) | 6.767 (14.468) | 0.640 ^—^ | 4.483 (2.100) | 0.033* ^a,b,c,d,g^ | 20.718 (25.783) | 0.424 ^—^ | 7.567 (4.125) | 0.068 ^a^ | -4.276 (16.404) | 0.795 ^—^ | 3.179 (2.386) | 0.184 ^a,c,d^ |
| Trail Making (B/A) | 0.021 (0.317) | 0.948 ^—^ | 0.047 (0.056) | 0.397 ^a,c,d^ | 0.052 (0.493) | 0.917 ^—^ | 0.133 (0.107) | 0.212 ^a^ | -0.021 (0.418) | 0.961 ^—^ | -0.008 (0.066) | 0.904 ^c,d^ |
| Verbal Fluency, Animals (Total) | -1.626 (1.563) | 0.299 ^—^ | -0.211 (0.208) | 0.311 ^a,b,c^ | 0.674 (2.346) | 0.775 ^—^ | -0.204 (0.418) | 0.625 ^a,c^ | -3.956 (2.142) | 0.067 ^—^ | -0.091 (0.233) | 0.696 ^a,c^ |
| Verbal Fluency, F+A+S (Total) | -2.352 (4.029) | 0.560 ^—^ | 0.071 (0.398) | 0.859 ^b,c,d^ | 1.916 (5.942) | 0.748 ^—^ | -0.113 (0.725) | 0.876 ^—^ | -6.041 (5.561) | 0.279 ^—^ | 0.240 (0.490) | 0.625 ^c,d^ |

**Supplemental Table 9.** Associations between baseline GFAP and longitudinal cognitive domain

scores in a) the whole ADNI cohort, b) males only, and c) females only. Output of linear mixed effects models. Each model was adjusted for age at baseline, time since baseline, race, education, and APOE4 status (sex was additionally adjusted for in the whole-cohort model). Significant associations are indicated with asterisks (p<0.05*, p<0.01**, p<0.001***).

| **Outcome (longitudinal)** | **Baseline GFAP, β (SE)** | **Baseline GFAP, p-value** | **Significant Covariates** |
| --- | --- | --- | --- |
| **a) Whole ADNI Cohort (N=370 participants across 1,191 total visits)** | | | |
| AVLT, Total Learned | -0.809 (2.519) | 0.748 | age at baseline, sex, education, time since baseline |
| AVLT, Delayed Recall | -0.395 (0.939) | 0.674 | age at baseline, sex, race, education, time since baseline |
| Trail Making A (Time) | 2.333 (2.669) | 0.383 | age at baseline, sex, race, time since baseline |
| Trail Making B (Time) | 25.184 (9.293) | 0.007** | age at baseline, sex, race, education, time since baseline |
| Verbal Fluency, Animals (Total) | -3.540 (1.135) | 0.002** | age at baseline, sex, race, education, time since baseline |
| Verbal Fluency, F (Total) | -1.682 (1.115) | 0.132 | education |
| **b) Males Only (N=153 participants across 528 total visits)** | | | |
| AVLT, Total Learned | -3.373 (3.858) | 0.383 | age at baseline, education, time since baseline |
| AVLT, Delayed Recall | -0.943 (1.440) | 0.514 | age at baseline, time since baseline |
| Trail Making A (Time) | 0.819 (4.626) | 0.860 | age at baseline, race, time since baseline |
| Trail Making B (Time) | 12.000 (15.778) | 0.448 | age at baseline, race, time since baseline |
| Verbal Fluency, Animals (Total) | -4.785 (1.755) | 0.007** | age at baseline, race, education, time since baseline |
| Verbal Fluency, F (Total) | -3.537 (1.732) | 0.043* | education |
| **c) Females Only (N=217 participants across 663 total visits)** | | | |
| AVLT, Total Learned | 0.649 (3.348) | 0.847 | age at baseline, race, education, time since baseline |
| AVLT, Delayed Recall | -0.077 (1.244) | 0.950 | age at baseline, race, education, time since baseline |
| Trail Making A (Time) | 2.768 (3.160) | 0.382 | age at baseline, race, time since baseline |
| Trail Making B (Time) | 35.275 (11.324) | 0.002** | age at baseline, race, time since baseline |
| Verbal Fluency, Animals (Total) | -2.701 (1.501) | 0.073 | age at baseline, race, education, time since baseline |
| Verbal Fluency, F (Total) | -0.427 (1.476) | 0.773 | education |

**Supplemental Table 10.** Associations between baseline GFAP and longitudinal MMSE in the ADNI cohort, including three-way interactions between baseline GFAP, sex, and time since baseline. Output of linear mixed effects models. Females are the reference group for sex interactions. Each model was adjusted for age at baseline, race, education, and APOE4 status, with time since baseline included as part of the interaction model. Significant associations are indicated with asterisks (p<0.05*, p<0.01**, p<0.001***).

| **Outcome (longitudinal)** | **GFAP** | | **GFAP × Time** | | **GFAP × Sex** | | **Sex × Time** | | **GFAP × Sex × Time** | | **Significant Covariates** |
| --- | --- | --- | --- | --- | --- | --- | --- | --- | --- | --- | --- |
|  | **β (SE)** | **p-value** | **β (SE)** | **p-value** | **β (SE)** | **p-value** | **β (SE)** | **p-value** | **β (SE)** | **p-value** |  |
| MMSE | -0.391 (0.488) | 0.423 | -0.099 (0.085) | 0.244 | -0.207 (0.683) | 0.762 | 0.329 (0.248) | 0.184 | -0.103 (0.116) | 0.375 | age at baseline, education |

**Supplemental Table 10b**. Associations between baseline GFAP and longitudinal MMSE for the interaction between baseline GFAP and time, in the full ADNI cohort, males only, and females only. Output of linear mixed effects models. Full-cohort models were adjusted for age at baseline, race, education, and APOE4 status; male- and female-only models were adjusted for age at baseline, race, education, and APOE4 status. Significant associations are indicated with asterisks (p<0.05*, p<0.01**, p<0.001***). Superscript letters on the GFAP × Time p-value indicate covariates significantly associated with the outcome in that model: a = age at baseline, b = sex, c = race, d = education, e = APOE4, f = time since baseline, g = time since baseline:GFAP interaction; — = no significant covariates.

| **Outcome (longitudinal)** | **Full Cohort** | | | | **Males** | | | | **Females** | | | |
| --- | --- | --- | --- | --- | --- | --- | --- | --- | --- | --- | --- | --- |
|  | **GFAP β (SE)** | **GFAP p-value** | **GFAP × Time β (SE)** | **GFAP × Time p-value** | **GFAP β (SE)** | **GFAP p-value** | **GFAP × Time β (SE)** | **GFAP × Time p-value** | **GFAP β (SE)** | **GFAP p-value** | **GFAP × Time β (SE)** | **GFAP × Time p-value** |
| MMSE | -0.360 (0.371) | 0.333 ^—^ | -0.190 (0.057) | 0.001*** ^a,b,d,g^ | -0.552 (0.575) | 0.337 ^—^ | -0.212 (0.079) | 0.008*** ^a,d,g^ | -0.449 (0.488) | 0.358 ^—^ | -0.094 (0.084) | 0.266 ^d^ |

**Supplemental Table 10c**. Associations between baseline GFAP and longitudinal cognition in the ADNI cohort, including three-way interactions between baseline GFAP, sex, and time since baseline. Output of linear mixed effects models. Females are the reference group for sex interactions. Each model was adjusted for age at baseline, race, education, and APOE4 status, with time since baseline included as part of the interaction model. Significant associations are indicated with asterisks (p<0.05*, p<0.01**, p<0.001***).

| **Outcome (longitudinal)** | **GFAP** | | **GFAP × Time** | | **GFAP × Sex** | | **Sex × Time** | | **GFAP × Sex × Time** | | **Significant Covariates** |
| --- | --- | --- | --- | --- | --- | --- | --- | --- | --- | --- | --- |
|  | **β (SE)** | **p-value** | **β (SE)** | **p-value** | **β (SE)** | **p-value** | **β (SE)** | **p-value** | **β (SE)** | **p-value** |  |
| AVLT, Total Learned | 1.826 (3.259) | 0.575 | -0.438 (0.478) | 0.360 | -4.025 (4.805) | 0.403 | 0.172 (1.500) | 0.909 | -0.079 (0.709) | 0.912 | age at baseline, education |
| AVLT, Delayed Recall | 0.385 (1.235) | 0.755 | -0.151 (0.202) | 0.457 | -1.467 (1.819) | 0.420 | -0.302 (0.634) | 0.634 | 0.154 (0.300) | 0.609 | age at baseline, race, education |
| Trail Making A (Time) | -2.081 (3.588) | 0.562 | 1.402 (0.670) | 0.037* | 8.723 (5.282) | 0.099 | 5.057 (2.097) | 0.016* | -2.410 (0.991) | 0.015* | age at baseline, race |
| Trail Making B (Time) | 4.802 (12.479) | 0.701 | 9.790 (2.284) | <0.001*** | 9.362 (18.373) | 0.611 | 12.963 (7.152) | 0.070 | -6.050 (3.380) | 0.074 | age at baseline, race, education |
| Trail Making (B-A) | 6.844 (10.782) | 0.526 | 8.396 (2.195) | <0.001*** | 0.730 (15.859) | 0.963 | 8.570 (6.865) | 0.212 | -3.942 (3.243) | 0.224 | age at baseline, race, education |
| Trail Making (B/A) | 0.362 (0.278) | 0.192 | 0.099 (0.067) | 0.138 | -0.345 (0.407) | 0.398 | -0.053 (0.207) | 0.800 | 0.031 (0.098) | 0.755 | age at baseline, education |
| Verbal Fluency, Animals (Total) | -0.303 (1.519) | 0.842 | -0.971 (0.280) | 0.001*** | -4.754 (2.237) | 0.034* | -1.934 (0.875) | 0.027* | 0.927 (0.414) | 0.025* | age at baseline, race, education |
| Verbal Fluency, F (Total) | -0.352 (1.461) | 0.810 | -0.247 (0.237) | 0.299 | -3.220 (2.153) | 0.136 | -1.111 (0.744) | 0.136 | 0.547 (0.352) | 0.120 | education |

**Supplemental Table 10d**. Associations between baseline GFAP and longitudinal cognition for the interaction between baseline GFAP and time, in the full ADNI cohort, males only, and females only. Output of linear mixed effects models. Full-cohort models were adjusted for age at baseline, race, education, and APOE4 status; male- and female-only models were adjusted for age at baseline, race, education, and APOE4 status. Significant associations are indicated with asterisks (p<0.05*, p<0.01**, p<0.001***). Superscript letters on the GFAP × Time p-value indicate covariates significantly associated with the outcome in that model: a = age at baseline, b = sex, c = race, d = education, e = APOE4, f = time since baseline, g = time since baseline:GFAP interaction; — = no significant covariates.

| **Outcome (longitudinal)** | **Full Cohort** | | | | **Males** | | | | **Females** | | | |
| --- | --- | --- | --- | --- | --- | --- | --- | --- | --- | --- | --- | --- |
|  | **GFAP β (SE)** | **GFAP p-value** | **GFAP × Time β (SE)** | **GFAP × Time p-value** | **GFAP β (SE)** | **GFAP p-value** | **GFAP × Time β (SE)** | **GFAP × Time p-value** | **GFAP β (SE)** | **GFAP p-value** | **GFAP × Time β (SE)** | **GFAP × Time p-value** |
| AVLT, Total Learned | 0.269 (2.640) | 0.919 ^—^ | -0.479 (0.349) | 0.171 ^a,b,d^ | -2.219 (4.035) | 0.583 ^—^ | -0.506 (0.506) | 0.318 ^a,d^ | 1.602 (3.518) | 0.649 ^—^ | -0.432 (0.491) | 0.380 ^a,c,d^ |
| AVLT, Delayed Recall | -0.195 (0.997) | 0.845 ^—^ | -0.088 (0.148) | 0.550 ^a,b,c,d^ | -0.954 (1.513) | 0.529 ^—^ | 0.005 (0.202) | 0.980 ^a^ | 0.247 (1.334) | 0.853 ^—^ | -0.146 (0.216) | 0.499 ^a,c,d^ |
| Trail Making A (Time) | 1.621 (2.895) | 0.576 ^—^ | 0.311 (0.490) | 0.525 ^a,b,c^ | 3.107 (5.025) | 0.537 ^—^ | -0.980 (0.850) | 0.250 ^a,c^ | -0.232 (3.386) | 0.945 ^—^ | 1.347 (0.562) | 0.017* ^a,c,g^ |
| Trail Making B (Time) | 9.324 (10.059) | 0.354 ^—^ | 6.955 (1.669) | <0.001*** ^a,b,c,d,f,g^ | 2.490 (16.818) | 0.882 ^—^ | 4.135 (2.463) | 0.094 ^a,c^ | 13.455 (12.421) | 0.280 ^—^ | 9.689 (2.310) | <0.001*** ^a,c,f,g^ |
| Trail Making (B-A) | 7.752 (8.648) | 0.370 ^—^ | 6.498 (1.601) | <0.001*** ^a,c,d,f,g^ | -0.892 (14.209) | 0.950 ^—^ | 4.871 (2.328) | 0.037* ^a,c,g^ | 14.205 (10.852) | 0.191 ^—^ | 8.243 (2.245) | <0.001*** ^a,c,d,f,g^ |
| Trail Making (B/A) | 0.229 (0.220) | 0.299 ^—^ | 0.109 (0.048) | 0.025* ^a,d,g^ | -0.083 (0.338) | 0.806 ^—^ | 0.138 (0.069) | 0.044* ^a,d,g^ | 0.498 (0.292) | 0.089 ^—^ | 0.092 (0.069) | 0.183 ^c,d^ |
| Verbal Fluency, Animals (Total) | -2.268 (1.227) | 0.065 ^—^ | -0.557 (0.204) | 0.007** ^a,b,c,d,f,g^ | -4.667 (1.873) | 0.014* ^—^ | -0.051 (0.282) | 0.857 ^a,c,d^ | -0.521 (1.636) | 0.750 ^—^ | -0.972 (0.296) | 0.001*** ^a,c,d,f,g^ |
| Verbal Fluency, F (Total) | -1.654 (1.183) | 0.163 ^—^ | -0.012 (0.174) | 0.944 ^d^ | -4.232 (1.812) | 0.021* ^—^ | 0.305 (0.243) | 0.211 ^d,e^ | 0.132 (1.576) | 0.934 ^—^ | -0.251 (0.249) | 0.314 ^d^ |

**Supplemental Table 11.** Associations between longitudinal GFAP and longitudinal BBMs, MMSE, and cognitive domain scores in a) the full MERI cohort, b) males only, and c) females. Output of linear mixed effects models. N and visit totals are reported once per outcome domain, as sample composition differs slightly by domain due to missing data. Each model was adjusted for age at visit, time since baseline, race, education, and APOE4 status (sex was additionally adjusted for in the full-cohort models). Significant associations are indicated with asterisks (p<0.05*, p<0.01**, p<0.001***).

| **Outcome (longitudinal)** | **Longitudinal GFAP, β (SE)** | **Longitudinal GFAP, p-value** | **Significant Covariates** |
| --- | --- | --- | --- |
| **a) Full Cohort** | | | |
| ***Blood-Based Biomarkers (N=210 participants across 605 total visits)*** | | | |
| Aβ40 | 45.147 (6.839) | <0.001*** | age at visit |
| Aβ42 | 2.067 (0.428) | <0.001*** | education, APOE4 |
| Aβ42/40 | -0.007 (0.002) | 0.001** | APOE4 |
| p-Tau231 | 0.228 (0.038) | <0.001*** | age at visit, APOE4 |
| NfL | 0.535 (0.036) | <0.001*** | age at visit |
| ***MMSE (N=212 participants across 612 total visits)*** | | | |
| MMSE | -1.142 (0.504) | 0.024* | age at visit |
| ***Cognitive Testing (N=209 participants across 600 total visits)*** | | | |
| AVLT, Total Learned | -9.430 (2.326) | <0.001*** | age at visit |
| AVLT, Delayed Recall | -2.378 (0.809) | 0.003** | age at visit |
| Trail Making A (Time) | 4.183 (3.110) | 0.179 | age at visit, APOE4 |
| Trail Making B (Time) | 19.512 (11.971) | 0.104 | age at visit, education |
| Trail Making (B-A) | 17.968 (10.848) | 0.098 | age at visit, education |
| Trail Making (B/A) | 0.330 (0.268) | 0.220 | age at visit, education |
| Verbal Fluency, Animals (Total) | -3.014 (1.206) | 0.013* | age at visit |
| Verbal Fluency, F+A+S (Total) | -1.688 (2.641) | 0.523 | education |
| **b) Males Only** | | | |
| ***Blood-Based Biomarkers (N=86 participants across 256 total visits)*** | | | |
| Aβ40 | 50.739 (10.996) | <0.001*** | No significant covariates |
| Aβ42 | 3.188 (0.644) | <0.001*** | APOE4 |
| Aβ42/40 | -0.004 (0.003) | 0.178 | APOE4 |
| p-Tau231 | 0.204 (0.053) | <0.001*** | age at visit |
| NfL | 0.627 (0.059) | <0.001*** | age at visit |
| ***MMSE (N=87 participants across 266 total visits)*** | | | |
| MMSE | -1.604 (0.718) | 0.027* | age at visit |
| ***Cognitive Testing (N=85 participants across 254 total visits)*** | | | |
| AVLT, Total Learned | -11.657 (3.390) | 0.001** | No significant covariates |
| AVLT, Delayed Recall | -2.653 (1.180) | 0.025* | No significant covariates |
| Trail Making A (Time) | 1.312 (4.925) | 0.790 | age at visit |
| Trail Making B (Time) | 43.901 (19.062) | 0.022* | age at visit |
| Trail Making (B-A) | 41.989 (17.428) | 0.017* | age at visit |
| Trail Making (B/A) | 0.644 (0.439) | 0.145 | age at visit |
| Verbal Fluency, Animals (Total) | -0.005 (1.947) | 0.998 | age at visit |
| Verbal Fluency, F+A+S (Total) | 0.780 (4.093) | 0.849 | No significant covariates |
| **c) Females Only** | | | |
| ***Blood-Based Biomarkers (N=124 participants across 349 total visits)*** | | | |
| Aβ40 | 41.804 (8.677) | <0.001*** | No significant covariates |
| Aβ42 | 1.005 (0.575) | 0.082 | age at visit, APOE4 |
| Aβ42/40 | -0.009 (0.003) | 0.003** | APOE4 |
| p-Tau231 | 0.252 (0.053) | <0.001*** | age at visit, APOE4 |
| NfL | 0.448 (0.045) | <0.001*** | age at visit |
| ***MMSE (N=125 participants across 360 total visits)*** | | | |
| MMSE | -0.665 (0.723) | 0.358 | age at visit |
| ***Cognitive Testing (N=124 participants across 346 total visits)*** | | | |
| AVLT, Total Learned | -6.869 (3.211) | 0.033* | age at visit |
| AVLT, Delayed Recall | -1.840 (1.109) | 0.098 | age at visit |
| Trail Making A (Time) | 4.908 (3.968) | 0.217 | age at visit, APOE4 |
| Trail Making B (Time) | -2.141 (15.125) | 0.888 | age at visit, education |
| Trail Making (B-A) | -1.467 (13.523) | 0.914 | age at visit, education |
| Trail Making (B/A) | 0.063 (0.334) | 0.850 | education, APOE4 |
| Verbal Fluency, Animals (Total) | -5.411 (1.497) | <0.001*** | age at visit |
| Verbal Fluency, F+A+S (Total) | -3.029 (3.462) | 0.382 | education |

**Supplemental Table 12.** Associations between Rate of Change-GFAP and Rate of Change BBMs, a) the MERI cohort and b) the ADNI cohort, each in the full cohort, males only, and females only (replaces former Supplemental Tables 17 and 18). Output of linear mixed effects models. N represents unique participants; timepoints shown in parentheses. Full-cohort models were additionally adjusted for sex. Significant associations are indicated with asterisks (p<0.05*, p<0.01**, p<0.001***). Superscript letters indicate covariates significantly associated with the outcome in that model: a = age, b = time between visits; — = no significant covariates.

| **Outcome (Rate of Change)** | **Full Cohort** | | | **Males** | | | **Females** | | |
| --- | --- | --- | --- | --- | --- | --- | --- | --- | --- |
|  | **N (timepoints)** | **GFAP Rate of Change, β (SE)** | **p-value** | **N (timepoints)** | **GFAP Rate of Change, β (SE)** | **p-value** | **N (timepoints)** | **GFAP Rate of Change, β (SE)** | **p-value** |
| **a) MERI Cohort** | | | | | | | | | |
| Aβ40 | 199 (388) | 73.648 (12.549) | <0.001*** ^—^ | 79 (167) | 113.229 (21.010) | <0.001*** ^—^ | 120 (221) | 36.978 (14.836) | 0.013* ^—^ |
| Aβ42 | 199 (388) | 3.487 (0.691) | <0.001*** ^—^ | 79 (167) | 5.628 (1.122) | <0.001*** ^—^ | 120 (221) | 1.465 (0.851) | 0.086 ^—^ |
| Aβ42/40 | 199 (388) | -0.011 (0.003) | <0.001*** ^—^ | 79 (167) | -0.017 (0.005) | <0.001*** ^—^ | 120 (221) | -0.005 (0.003) | 0.084 ^—^ |
| p-Tau231 | 199 (388) | 0.136 (0.054) | 0.012* ^—^ | 79 (167) | 0.043 (0.072) | 0.554 ^—^ | 120 (221) | 0.232 (0.079) | 0.004** ^—^ |
| NfL | 199 (388) | 0.490 (0.042) | <0.001*** ^—^ | 79 (167) | 0.523 (0.073) | <0.001*** ^—^ | 120 (221) | 0.457 (0.048) | <0.001*** ^—^ |
| **b) ADNI Cohort** | | | | | | | | | |
| Aβ40 | 223 (254) | 0.136 (0.024) | <0.001*** ^—^ | 98 (110) | 0.077 (0.049) | 0.117 ^—^ | 125 (144) | 0.170 (0.022) | <0.001 ^a,b^ |
| Aβ42 | 223 (254) | 0.198 (0.022) | <0.001*** ^—^ | 98 (110) | 0.189 (0.045) | <0.001*** ^—^ | 125 (144) | 0.204 (0.023) | <0.001 ^b^ |
| Aβ42/40 | 223 (254) | 0.005 (0.001) | <0.001*** ^—^ | 98 (110) | 0.009 (0.002) | <0.001*** ^—^ | 125 (144) | 0.003 (0.001) | 0.032 ^b^ |
| p-Tau217 | 223 (254) | 0.098 (0.012) | <0.001*** ^b^ | 98 (110) | 0.127 (0.022) | <0.001*** ^b^ | 125 (144) | 0.082 (0.013) | <0.001 ^—^ |
| NfL | 223 (254) | 0.421 (0.036) | <0.001*** ^a^ | 98 (110) | 0.332 (0.053) | <0.001*** ^a^ | 125 (144) | 0.468 (0.048) | <0.001 ^—^ |

**Supplemental Table 13**. Associations between baseline GFAP and longitudinal blood-based biomarkers, global cognition (MMSE), and domain-specific cognitive testing for the interaction between baseline GFAP and age at baseline in MERI participants. Output of linear mixed-effects models. Each model was adjusted for sex, race, education, APOE4 status, and time since baseline. Significant associations are indicated with asterisks (p<0.05*, p<0.01**, p<0.001***).

| **Outcome (longitudinal)** | **N** | **N (unique)** | **GFAP (baseline), β (SE)** | **GFAP (baseline), p-value** | **GFAP × Age, β (SE)** | **GFAP × Age, p-value** | **Significant Covariates** |
| --- | --- | --- | --- | --- | --- | --- | --- |
| **Blood-Based Biomarkers** | | | | | | | |
| Aβ40 | 605 | 210 | -10.059 (61.562) | 0.870 | 0.455 (0.859) | 0.597 | time since baseline |
| Aβ42 | 605 | 210 | -0.651 (4.062) | 0.873 | 0.015 (0.057) | 0.792 | education, APOE4, time since baseline |
| Aβ42/40 | 605 | 210 | -0.009 (0.026) | 0.727 | 0.000 (0.000) | 0.960 | APOE4 |
| p-Tau231 | 605 | 210 | -0.432 (0.377) | 0.253 | 0.008 (0.005) | 0.110 | sex, APOE4, time since baseline |
| NfL | 605 | 210 | 0.543 (0.432) | 0.210 | -0.003 (0.006) | 0.665 | race, time since baseline |
| **MMSE** | | | | | | | |
| MMSE | 765 | 245 | 5.770 (4.203) | 0.171 | -0.109 (0.059) | 0.065 | sex, race, education, time since baseline |
| **Cognitive Testing** | | | | | | | |
| AVLT, Total Learned | 735 | 242 | 40.344 (24.986) | 0.108 | -0.699 (0.349) | 0.046* | sex, race, education, time since baseline |
| AVLT, Delayed Recall | 735 | 242 | 8.571 (9.044) | 0.344 | -0.175 (0.126) | 0.166 | sex, time since baseline |
| Trail Making A, Time | 735 | 242 | -36.852 (33.500) | 0.273 | 0.578 (0.468) | 0.218 | sex, race, time since baseline |
| Trail Making B, Time | 735 | 242 | -120.540 (127.413) | 0.345 | 2.003 (1.778) | 0.261 | sex, race, education, time since baseline |
| Trail Making B minus A | 735 | 242 | -84.163 (104.406) | 0.421 | 1.424 (1.456) | 0.329 | sex, race, education, time since baseline |
| Trail Making B/A Ratio | 735 | 242 | -0.317 (2.228) | 0.887 | 0.006 (0.031) | 0.838 | race, education, time since baseline |
| Verbal Fluency, Animals | 735 | 242 | 1.035 (11.464) | 0.928 | -0.044 (0.160) | 0.783 | sex, race, time since baseline |
| Verbal Fluency, FAS | 735 | 242 | -3.325 (30.142) | 0.912 | 0.016 (0.421) | 0.970 | sex, race, education, time since baseline |

**Supplemental Table 14**. Associations between baseline GFAP and longitudinal blood-based biomarkers, global cognition (MMSE), and domain-specific cognitive testing for the interaction between baseline GFAP and APOE4 status in MERI participants. Output of linear mixed-effects models. Each model was adjusted for age, sex, race, education, and time since baseline. Significant associations are indicated with asterisks (p<0.05*, p<0.01**, p<0.001***).

| **Outcome (longitudinal)** | **N** | **N (unique)** | **GFAP (baseline), β (SE)** | **GFAP (baseline), p-value** | **GFAP × APOE4, β (SE)** | **GFAP × APOE4, p-value** | **Significant Covariates** |
| --- | --- | --- | --- | --- | --- | --- | --- |
| **Blood-Based Biomarkers** | | | | | | | |
| Aβ40 | 605 | 210 | 21.058 (8.877) | 0.019* | 4.465 (15.270) | 0.770 | age at baseline, time since baseline |
| Aβ42 | 605 | 210 | 0.577 (0.590) | 0.330 | -0.588 (1.011) | 0.561 | age at baseline, education, time since baseline |
| Aβ42/40 | 605 | 210 | -0.006 (0.004) | 0.110 | -0.006 (0.006) | 0.368 | No significant covariates |
| p-Tau231 | 605 | 210 | 0.154 (0.055) | 0.006** | 0.051 (0.094) | 0.590 | age at baseline, sex, time since baseline |
| NfL | 605 | 210 | 0.369 (0.063) | <0.001*** | -0.044 (0.108) | 0.682 | age at baseline, race, time since baseline |
| **MMSE** | | | | | | | |
| MMSE | 765 | 245 | -1.769 (0.644) | 0.007** | -0.682 (1.063) | 0.522 | age at baseline, sex, race, education, time since baseline |
| **Cognitive Testing** | | | | | | | |
| AVLT, Total Learned | 735 | 242 | -6.684 (3.744) | 0.076 | -8.699 (6.101) | 0.155 | age at baseline, sex, race, education, time since baseline |
| AVLT, Delayed Recall | 735 | 242 | -3.012 (1.352) | 0.027* | -2.909 (2.203) | 0.188 | age at baseline, sex, time since baseline |
| Trail Making A, Time | 735 | 242 | 1.643 (5.012) | 0.743 | 8.504 (8.166) | 0.299 | age at baseline, sex, race, time since baseline |
| Trail Making B, Time | 735 | 242 | 20.969 (19.055) | 0.272 | 2.815 (31.043) | 0.928 | age at baseline, sex, race, education, time since baseline |
| Trail Making B minus A | 735 | 242 | 18.369 (15.532) | 0.238 | -4.379 (25.307) | 0.863 | age at baseline, sex, race, education, time since baseline |
| Trail Making B/A Ratio | 735 | 242 | 0.235 (0.327) | 0.473 | -0.334 (0.534) | 0.532 | age at baseline, race, education, time since baseline |
| Verbal Fluency, Animals | 735 | 242 | -1.356 (1.704) | 0.427 | -2.505 (2.776) | 0.368 | age at baseline, sex, race, time since baseline |
| Verbal Fluency, FAS | 735 | 242 | 1.063 (4.487) | 0.813 | -10.944 (7.312) | 0.136 | sex, race, education, time since baseline |

**Supplemental Table 15.** Associations between baseline GFAP and longitudinal BBMs, global cognition (MMSE), and domain-specific cognitive testing for the interaction between baseline GFAP and age at baseline in ADNI participants. Output of linear mixed-effects models. Each model was adjusted for sex, race, education, APOE4 status, and time since baseline. Significant associations are indicated with asterisks (p<0.05*, p<0.01**, p<0.001***).

| **Outcome (longitudinal)** | **N** | **N (unique)** | **GFAP (baseline), β (SE)** | **GFAP (baseline), p-value** | **GFAP × Age, β (SE)** | **GFAP × Age, p-value** | **Significant Covariates** |
| --- | --- | --- | --- | --- | --- | --- | --- |
| **Blood-Based Biomarkers** | | | | | | | |
| Aβ40 | 675 | 294 | 0.209 (0.222) | 0.347 | -0.002 (0.003) | 0.462 | education, APOE4, time since baseline |
| Aβ42 | 675 | 294 | 0.115 (0.199) | 0.563 | -0.001 (0.003) | 0.594 | education, APOE4, time since baseline |
| Aβ42/40 | 675 | 294 | -0.005 (0.017) | 0.769 | 0.000 (0.000) | 0.895 | APOE4, time since baseline |
| p-Tau217 | 675 | 294 | 0.087 (0.134) | 0.520 | 0.000 (0.002) | 0.968 | race, APOE4, time since baseline |
| NfL | 548 | 294 | -0.308 (0.391) | 0.430 | 0.009 (0.005) | 0.107 | race, APOE4, time since baseline |
| **MMSE** | | | | | | | |
| MMSE | 1,905 | 447 | -2.473 (3.340) | 0.459 | 0.021 (0.046) | 0.64 | sex, education, time since baseline |
| **Cognitive Testing** | | | | | | | |
| AVLT, Total Learned | 1,191 | 370 | -18.849 (21.963) | 0.391 | 0.252 (0.305) | 0.409 | sex, education, time since baseline |
| AVLT, Delayed Recall | 1,191 | 370 | -3.818 (8.222) | 0.643 | 0.048 (0.114) | 0.675 | sex, race, education, time since baseline |
| Trail Making A, Time | 1,191 | 370 | 5.764 (23.492) | 0.806 | -0.048 (0.326) | 0.883 | sex, race, time since baseline |
| Trail Making B, Time | 1,191 | 370 | 0.404 (81.776) | 0.996 | 0.346 (1.136) | 0.761 | sex, race, education, time since baseline |
| Trail Making B minus A | 1,191 | 370 | 0.591 (69.071) | 0.993 | 0.310 (0.959) | 0.747 | race, education, time since baseline |
| Trail Making B/A Ratio | 1,191 | 370 | -1.273 (1.682) | 0.450 | 0.025 (0.023) | 0.293 | education, time since baseline |
| Verbal Fluency, Animals | 1,191 | 370 | -19.703 (9.947) | 0.048* | 0.226 (0.138) | 0.103 | age at baseline, sex, race, education, time since baseline |
| Verbal Fluency, F | 1,191 | 370 | -9.910 (9.752) | 0.310 | 0.115 (0.135) | 0.396 | education |

**Supplemental Table 16**. Associations between baseline GFAP and longitudinal blood-based biomarkers, global cognition (MMSE), and domain-specific cognitive testing for the interaction between baseline GFAP and APOE4 status in ADNI participants. Output of linear mixed-effects models. Each model was adjusted for age, sex, race, education, and time since baseline. Significant associations are indicated with asterisks (p<0.05*, p<0.01**, p<0.001***).

| **Outcome (longitudinal)** | **N** | **N (unique)** | **GFAP (baseline), β (SE)** | **GFAP (baseline), p-value** | **GFAP × APOE4, β (SE)** | **GFAP × APOE4, p-value** | **Significant Covariates** |
| --- | --- | --- | --- | --- | --- | --- | --- |
| **Blood-Based Biomarkers** | | | | | | | |
| Aβ40 | 675 | 294 | 0.056 (0.028) | 0.046* | -0.026 (0.043) | 0.544 | age at baseline, education, time since baseline |
| Aβ42 | 675 | 294 | 0.019 (0.025) | 0.443 | -0.027 (0.039) | 0.486 | age at baseline, education, time since baseline |
| Aβ42/40 | 675 | 294 | -0.003 (0.002) | 0.238 | -0.001 (0.003) | 0.845 | time since baseline |
| p-Tau217 | 675 | 294 | 0.076 (0.017) | <0.001*** | 0.045 (0.026) | 0.084 | age at baseline, race, time since baseline |
| NfL | 548 | 294 | 0.311 (0.049) | <0.001*** | 0.025 (0.076) | 0.740 | age at baseline, race, time since baseline |
| **MMSE** | | | | | | | |
| MMSE | 1,905 | 447 | -0.851 (0.397) | 0.032* | -0.188 (0.620) | 0.761 | age at baseline, sex, education, time since baseline |
| **Cognitive Testing** | | | | | | | |
| AVLT, Total Learned | 1,191 | 370 | 1.463 (2.995) | 0.626 | -6.366 (4.555) | 0.163 | age at baseline, sex, education, time since baseline |
| AVLT, Delayed Recall | 1,191 | 370 | 0.636 (1.115) | 0.569 | -2.897 (1.698) | 0.089 | age at baseline, sex, race, education, time since baseline |
| Trail Making A, Time | 1,191 | 370 | -0.437 (3.171) | 0.891 | 7.802 (4.839) | 0.108 | age at baseline, sex, race, time since baseline |
| Trail Making B, Time | 1,191 | 370 | 27.558 (11.066) | 0.013* | -6.685 (16.882) | 0.692 | age at baseline, sex, race, education, time since baseline |
| Trail Making B minus A | 1,191 | 370 | 28.190 (9.266) | 0.003** | -15.422 (14.170) | 0.277 | age at baseline, race, education, time since baseline |
| Trail Making B/A Ratio | 1,191 | 370 | 0.763 (0.221) | 0.001*** | -0.787 (0.339) | 0.021* | age at baseline, education, APOE4, time since baseline |
| Verbal Fluency, Animals | 1,191 | 370 | -3.490 (1.352) | 0.010** | -0.139 (2.063) | 0.946 | age at baseline, sex, race, education, time since baseline |
| Verbal Fluency, F | 1,191 | 370 | -1.081 (1.328) | 0.416 | -1.689 (2.022) | 0.404 | education |

**Supplemental Table 17.** Cross-sectional mediation analyses of sex, GFAP, and BBM or cognitive in a) the MERI cohort and b) the ADNI cohort. Output of mediation models evaluating the indirect pathway from sex to each outcome through GFAP. Female was the reference group. N represents the outcome-specific complete-case sample size. Each mediator and outcome model was adjusted for age, race, education, and APOE4 status. ACME represents the average causal mediation effect (indirect effect) through GFAP, ADE represents the average direct effect of sex independent of GFAP, and total effect represents the overall association between sex and the outcome. Confidence intervals were estimated using 5,000 bootstrap resamples. Given the cross-sectional nature of these analyses, they assess statistical indirect associations but do not establish temporal or causal mediation. Significant associations are indicated with asterisks (p<0.05*, p<0.01**, p<0.001***).

| **Outcome** | **N** | **ACME, β [95% CI]** | **p** | **ADE, β [95% CI]** | **p** | **Total Effect, β [95% CI]** | **p** |
| --- | --- | --- | --- | --- | --- | --- | --- |
| **a) MERI Cohort** | | | | | | | |
| p-Tau231 | 583 | -0.029 [-0.045, -0.017] | <0.001*** | 0.074 [0.041, 0.108] | <0.001*** | 0.045 [0.012, 0.078] | 0.008** |
| Aβ40 | 581 | -3.579 [-5.721, -1.855] | <0.001*** | 14.284 [7.727, 21.257] | <0.001*** | 10.704 [4.585, 17.247] | <0.001*** |
| Aβ42 | 565 | -0.135 [-0.239, -0.048] | 0.002** | 0.553 [0.162, 0.948] | 0.005** | 0.418 [0.043, 0.794] | 0.026* |
| Aβ42/40 | 565 | 0.00085 [0.00035, 0.00150] | <0.001*** | -0.00189 [-0.00376, -0.00004] | 0.044* | -0.00104 [-0.00292, 0.00086] | 0.275 |
| NfL | 584 | -0.056 [-0.079, -0.036] | <0.001*** | 0.075 [0.042, 0.108] | <0.001*** | 0.019 [-0.018, 0.056] | 0.326 |
| MMSE | 584 | 0.125 [0.049, 0.218] | <0.001*** | -0.151 [-0.423, 0.115] | 0.266 | -0.026 [-0.289, 0.235] | 0.837 |
| AVLT, Total Learned | 582 | 0.950 [0.380, 1.635] | <0.001*** | -6.179 [-8.163, -4.216] | <0.001*** | -5.229 [-7.117, -3.343] | <0.001*** |
| AVLT, Delayed Recall | 581 | 0.332 [0.139, 0.563] | <0.001*** | -1.821 [-2.464, -1.161] | <0.001*** | -1.489 [-2.104, -0.842] | <0.001*** |
| Trail Making A, Time | 580 | -1.152 [-2.138, -0.379] | 0.004** | 4.767 [0.840, 9.215] | 0.016* | 3.615 [-0.322, 7.934] | 0.075 |
| Trail Making B, Time | 544 | -2.855 [-5.737, -0.214] | 0.031* | 6.261 [-4.354, 16.820] | 0.251 | 3.406 [-7.057, 13.978] | 0.517 |
| Trail Making B minus A | 544 | -1.962 [-4.399, 0.282] | 0.081 | 2.370 [-6.373, 11.465] | 0.586 | 0.408 [-8.308, 9.436] | 0.904 |
| Trail Making B/A Ratio | 544 | -0.010 [-0.055, 0.036] | 0.675 | -0.071 [-0.261, 0.112] | 0.449 | -0.081 [-0.267, 0.102] | 0.389 |
| Verbal Fluency, Animals | 579 | 0.420 [0.178, 0.713] | <0.001*** | -1.253 [-2.195, -0.317] | 0.010* | -0.834 [-1.757, 0.116] | 0.085 |
| Verbal Fluency, FAS | 581 | 0.580 [0.034, 1.208] | 0.037* | -3.917 [-6.368, -1.464] | 0.001** | -3.337 [-5.763, -0.906] | 0.006** |
| **b) ADNI Cohort** | | | | | | | |
| p-Tau217 | 707 | -0.007 [-0.011, -0.004] | <0.001*** | 0.009 [0.001, 0.017] | 0.021* | 0.002 [-0.006, 0.010] | 0.660 |
| Aβ40 | 707 | -0.005 [-0.010, -0.001] | 0.004** | -0.024 [-0.043, -0.006] | 0.008** | -0.029 [-0.047, -0.012] | 0.001** |
| Aβ42 | 707 | -0.001 [-0.005, 0.003] | 0.545 | -0.039 [-0.059, -0.019] | <0.001*** | -0.040 [-0.059, -0.021] | <0.001*** |
| Aβ42/40 | 707 | 0.00026 [0.00005, 0.00050] | 0.015* | -0.00062 [-0.00204, 0.00118] | 0.410 | -0.00036 [-0.00173, 0.00137] | 0.591 |
| NfL | 707 | -0.030 [-0.042, -0.019] | <0.001*** | 0.022 [0.000, 0.044] | 0.048* | -0.008 [-0.032, 0.016] | 0.516 |
| MMSE | 622 | 0.021 [-0.032, 0.072] | 0.413 | -0.481 [-0.701, -0.275] | <0.001*** | -0.460 [-0.682, -0.249] | <0.001*** |
| AVLT, Total Learned | 622 | -0.049 [-0.422, 0.319] | 0.778 | -6.510 [-8.224, -4.798] | <0.001*** | -6.560 [-8.194, -4.898] | <0.001*** |
| AVLT, Delayed Recall | 622 | 0.020 [-0.124, 0.171] | 0.772 | -2.301 [-2.974, -1.646] | <0.001*** | -2.280 [-2.937, -1.649] | <0.001*** |
| Trail Making A, Time | 622 | -0.311 [-0.756, 0.060] | 0.112 | 4.390 [2.513, 6.302] | <0.001*** | 4.078 [2.213, 5.921] | <0.001*** |
| Trail Making B, Time | 622 | -0.709 [-2.033, 0.510] | 0.261 | 2.015 [-4.454, 8.301] | 0.555 | 1.306 [-5.240, 7.659] | 0.700 |
| Trail Making B minus A | 622 | -0.398 [-1.554, 0.717] | 0.460 | -2.374 [-8.260, 3.526] | 0.422 | -2.773 [-8.726, 3.065] | 0.345 |
| Trail Making B/A Ratio | 622 | -0.004 [-0.039, 0.028] | 0.806 | -0.227 [-0.409, -0.059] | 0.007** | -0.231 [-0.411, -0.063] | 0.008** |
| Verbal Fluency, Animals | 622 | 0.079 [-0.091, 0.278] | 0.370 | -1.487 [-2.334, -0.618] | <0.001*** | -1.407 [-2.226, -0.581] | <0.001*** |
| Verbal Fluency, F | 622 | 0.034 [-0.131, 0.201] | 0.665 | -0.654 [-1.444, 0.149] | 0.115 | -0.620 [-1.384, 0.169] | 0.127 |
